# Adaptive laboratory evolution reveals general and specific chemical tolerance mechanisms and enhances biochemical production

**DOI:** 10.1101/634105

**Authors:** Rebecca M. Lennen, Kristian Jensen, Elsayed T. Mohammed, Sailesh Malla, Rosa A. Börner, Ksenia Chekina, Emre Özdemir, Ida Bonde, Anna Koza, Jérôme Maury, Lasse E. Pedersen, Lars Y. Schöning, Nikolaus Sonnenschein, Bernhard O. Palsson, Morten O.A. Sommer, Adam M. Feist, Alex T. Nielsen, Markus J. Herrgård

## Abstract

Tolerance to high product concentrations is a major barrier to achieving economically viable processes for bio-based chemical production. Chemical tolerance mechanisms are often unknown, thus their rational design is not achievable. To reveal unknown tolerance mechanisms we used an automated platform to evolve *Escherichia coli* to grow in previously toxic concentrations of 11 chemicals that have applications as polymer precursors, chemical intermediates, or biofuels. Re-sequencing of isolates from 88 independently evolved populations, reconstruction of mutations, and cross-compound tolerance profiling was employed to uncover general and specific tolerance mechanisms. We found that: 1) the broad tolerance of strains towards chemicals varied significantly depending on the chemical stress condition under which the strain was evolved; 2) the strains that acquired high levels of NaCl tolerance also became broadly tolerant to most chemicals; 3) genetic tolerance mechanisms included alterations in regulatory, cell wall, transcriptional and translational functions, as well as more chemical-specific mechanisms related to transport and metabolism; 4) using pre-tolerized starting strains can significantly enhance subsequent production of chemicals when a production pathway is inserted; and 5) only a subset of the evolved isolates showed improved production indicating that this approach is especially useful when a large number of independently evolved isolates are screened for production. We provide a comprehensive genotype-phenotype map based on identified mutations and growth phenotypes for 224 chemical tolerant strains.

## Introduction

Despite advances in synthetic and systems biology tools to engineer and study metabolism, developing microbial strains for commercial-level chemical production remains a challenge^1^. The stressful conditions that production strains encounter in large-scale industrial processes are one of the most significant hurdles for commercialization^2^. High concentrations of the compound that is being produced is one of the major stresses present in large-scale production conditions. Chemical stress can have inhibitory effects on the host organism, limiting the achievable titers and thereby the economic feasibility of the production process. These issues can be overcome by engineering a production strain that is tolerant to higher product titers, but this is rarely possible through rational engineering due to lack of knowledge about the molecular mechanisms of chemical toxicity or tolerance^3^. This requires either choosing a more robust, but potentially otherwise difficult to engineer production organism, or alternatively using non-rational approaches to engineer tolerance. These approaches include induced random mutagenesis, library screening, or adaptive laboratory evolution (ALE)^4^. ALE in particular has been successfully used to obtain strains that tolerate product chemicals^35^. In some cases the mechanisms of chemical tolerance in ALE-derived strains have been partially deciphered through resequencing and other omics approaches^6–8^, but in most cases the toxicity and tolerance mechanisms remain unknown. While some cases of ALE applied to product tolerance has resulted in strains that increase actual production of the target chemical^9^, in other cases no significant improvement in production has been observed^6,10^.

Here we take a broad approach to elucidating genetic mechanisms of chemical tolerance across a wide spectrum of chemicals enabled by automated ALE as well as systematic genomic and phenotypic analyses of the resulting large collection of evolved *Escherichia coli* strains. This approach allowed determining general features of chemical tolerance and building a comprehensive reference dataset for future tolerance studies. For two chemical products we also established that evolving for tolerance can significantly improve production, but that the degree of improvement depends on the specific genotype of the evolved strain.

## Results

We selected 11 chemical compounds representing a diversity of chemical categories with variable initial levels of toxicity to *E. coli* (Figure 1a). We chose the chemicals to include compounds with potential as bio-based products, cover multiple chemical compound classes, include chemicals belonging to the same compound classes, and to have compounds with high solubility and low volatility suitable for ALE. Two of the compounds (octanoate and n-butanol) had previously been used in ALE studies in *E. coli*^8,11^. For most of the compounds, there have been efforts to engineer improved production in *E. coli* (Supplementary Table 5).

We used an automated serial passaging platform to evolve eight independent populations of *E. coli* K-12 MG1655 to tolerate previously toxic levels of each of the 11 target chemicals, resulting in a total of 88 independently evolved populations. During the laboratory evolution process, we increased the chemical concentrations in a stepwise manner over approximately 800 generations. The starting and end concentrations that allowed population growth are shown in Figure 1b along with the overall percent increase over the course of evolution (60% - 400%). None of the evolved populations exhibited significant growth with the toxic compound as a sole carbon source, suggesting that they had not evolved the ability to degrade the compound. We tested ten isolates from each population for ability to grow in the final concentration of a chemical, and up to three isolates per population that grew robustly were selected for further characterization. This resulted in a total of 224 strains with evolved tolerance to one of the 11 chemicals. We subjected all strains to whole genome resequencing and cross-compound tolerance screening. In the cases of isobutyrate and 2,3-butanediol, we engineered production pathways into all genetically distinct isolates in order to determine if evolved product-tolerant strains exhibit increased production when the product is made endogenously. The overall workflow of the study is shown in Figure 1c.

**Figure 1:**
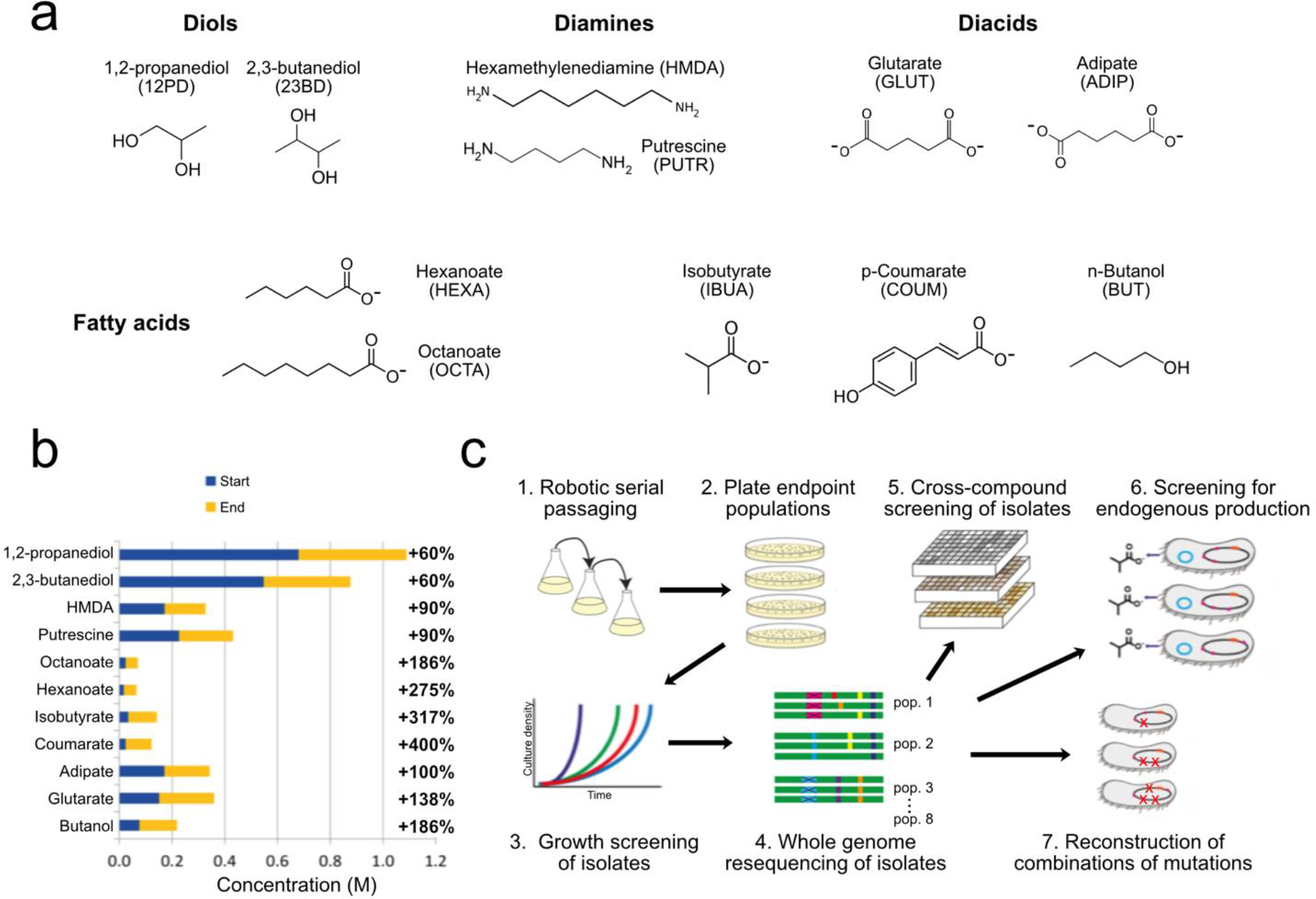
a) Chemicals selected for the study grouped by chemical category. b) Initial and final concentrations of the chemicals used during ALE and the percentage increase in tolerated concentration. c) Overall workflow of the study.

### Genome sequencing

The evolved isolates had a median number of sequence variants (excluding duplications) of 6, although a subset of the strains had more than 10 times this number of variants. This drastic difference was caused by a hypermutator phenotype in some strains, which possessed mutations in mismatch repair genes (e.g. *mutS*). Since the hypermutator strains were assumed to have accumulated mostly random neutral variants, they were not included in further analysis of sequence variants. The 1,2-propanediol condition was left out of this analysis as only three isolates from two out of eight populations were not hypermutators. The median number of variants among the remaining 189 strains was 5 and the numbers of variants for strains evolved in different conditions were similar (Figure 2a). A subset of strains, especially those evolved on isobutyrate and coumarate (Figure 2a), contained large duplications, To investigate which cellular functions were affected by the mutations, the functional annotations of all mutated genes were analyzed (Figure 2b). More than half of the variants affect genes with regulatory or transport functions, indicating that these gene classes play a major role in the evolution of tolerance.

We were able to determine potentially causal mutations by identifying genes that had mutations in isolates from many of the independently evolved populations for the same condition. In four conditions we identified genes that were mutated in all isolates from that condition: glutarate and adipate strains had *kgtP* mutations, isobutyrate strains had *pykF* mutations, and 2,3-butanediol strains had *relA* mutations. Furthermore, we observed mutations in a number of other genes in at least one strain from almost all populations (Table 1). There was limited overlap of mutated genes between the different evolution conditions (Supplementary Table 6). Only 12 genes had mutations in at least one isolate from four or more conditions: *hns, nagC, proV, pyrE, rpoA, rpoB, rpoC, rpsA, spoT, sspA, yeaR* and *yobF*. This list includes genes that likely have global regulatory effects (e.g. *rpsA, rpoABC*, *spoT*, *sspA* and *hns*), genes that are commonly found to be mutated in *E. coli* ALE studies^12^ (e.g. *pyrE*), and genes that have previously been found to be mutated in osmotolerance ALE studies (e.g. *nagC* and *proV*)^13^. In cases where the same gene was mutated in different evolution conditions, the specific mutations were usually distinct, indicating that the effects of the mutations may also be different (see Supplementary Figure 2 for RNA polymerase mutations).

**Table 1:** The five most commonly mutated genes for each condition. The numbers in parentheses denote the number of ALE populations in which mutations in the given gene were observed in at least one strain.

| HMDA | PUTR | 23BD | GLUT | ADIP | HEXA | OCTA | IBUA | COUM | BUT |
| --- | --- | --- | --- | --- | --- | --- | --- | --- | --- |
| *pyrE*<br>(4/6) | *mreB*<br>(5/7) | *metJ*<br>(7/7) | *kgtP*<br>(8/8) | *kgtP*<br>(7/7) | *rpoA*<br>(7/7) | *rpoC*<br>(3/6) | *pykF*<br>(8/8) | *rho*<br>(7/8) | *pyrE*<br>(7/8) |
| *proV*<br>(3/6) | *spoT*<br>(4/7) | *relA*<br>(7/7) | *spoT*<br>(7/8) | *ybjL*<br>(5/7) | *sapB*<br>(3/7) | *rpoA*<br>(2/6) | *rpoB*<br>(6/8) | *nadR*<br>(4/8) | *manY*<br>(7/8) |
| *nagC*<br>(3/6) | *rpoC*<br>(3/7) | *rpoC*<br>(5/7) | *rpoC*<br>(5/8) | *proV*<br>(4/7) | *mdtK*<br>(3/7) | *dusB*<br>(2/6) | *glyQ*<br>(2/8) | *pyrE*<br>(4/8) | *rob*<br>(7/8) |
| *ptsP*<br>(2/6) | *proV*<br>(3/7) | *nanK*<br>(5/7) | *nagC*<br>(4/8) | *pyrE*<br>(3/7) | *rpoC*<br>(3/7) | *gtrS*<br>(2/6) | *rpoC*<br>(2/8) | *manY*<br>(4/8) | *marC*<br>(6/8) |
| *ybeX*<br>(2/6) | *rpsA*<br>(3/7) | *purT*<br>(5/7) | *proV*<br>(3/8) | *sspA*<br>(3/7) | *ompC*<br>(2/7) | *mreB*<br>(2/6) | *rpoS*<br>(2/8) | *mprA*<br>(3/8) | *yobF*<br>(4/8) |

**Figure 2:**
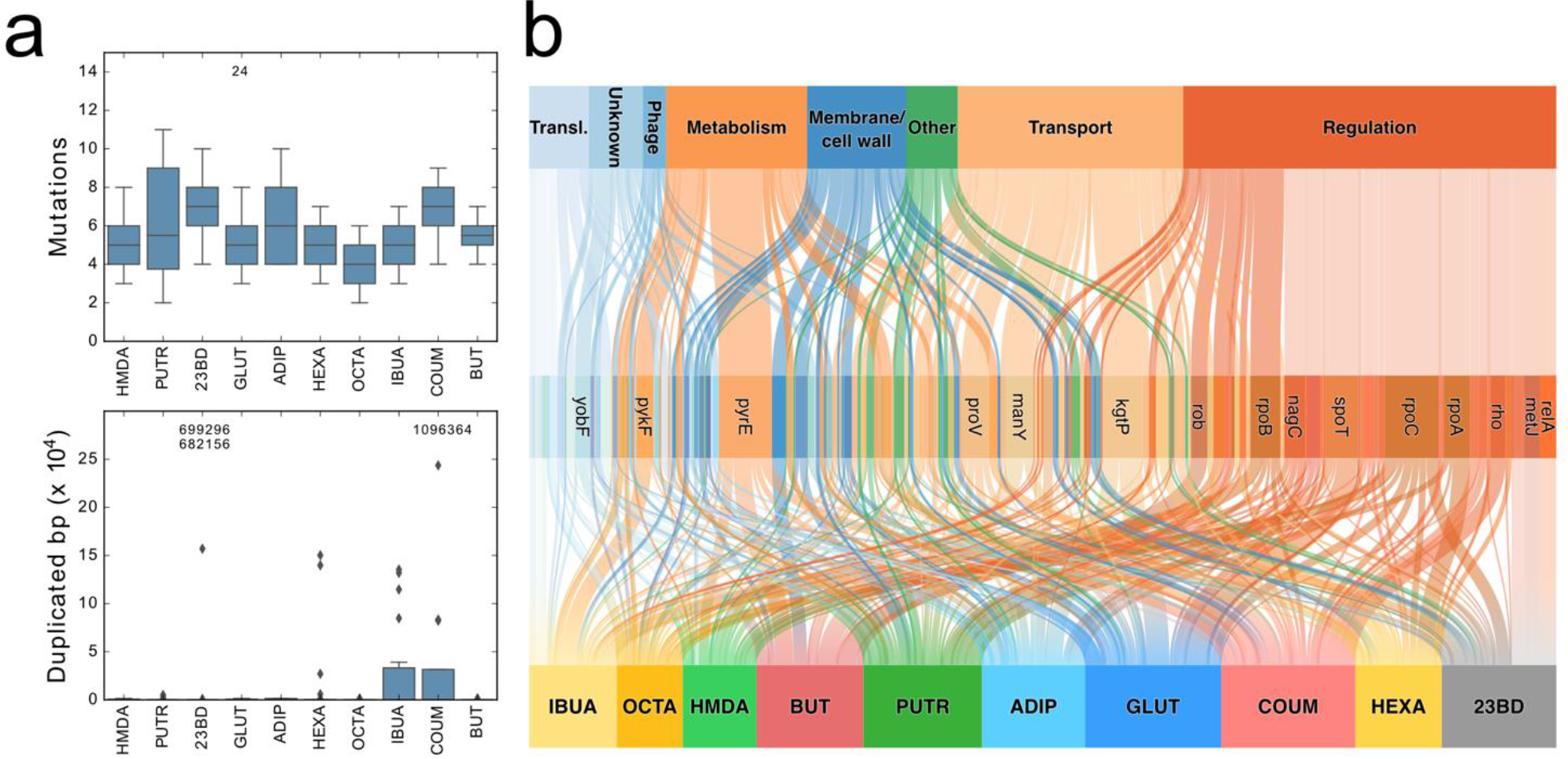
a) Boxplots showing the distributions of mutations per strain and duplication size per strain for each condition. The numbers above the boxes show the values of outliers not shown in the plots. b) Genetic variant landscape shown as a Sankey diagram. The chart shows an overview of the genes mutated in the different conditions and the functional classifications of these genes. The width of the lines is proportional to the number of strains in which a given gene was mutated.

### Cross-compound tolerance

In order to determine whether the strains had tolerance to a broad range of chemicals, we cultured all 224 isolates in the presence of moderately toxic levels of each of the 11 chemicals (Supplementary Table 1). We used the growth rate of a strain in a given condition relative to the wild-type strain as a measure of tolerance. Additionally, we grew the strains in M9 glucose to determine general growth improvements or tradeoffs, and in M9 glucose + 0.6 M NaCl to determine whether non-specific tolerance to high NaCl conditions (both osmotic and cation stress) was evolved. We found that strains evolved on diamines, diols and diacids were generally tolerant to the other chemical of the same functional class (Figure 3a). In contrast, strains evolved on either of the medium chain-length fatty acids (hexanoic or octanoic acid) were not tolerant to the other medium chain-length fatty acid. We also tested whether strains that were evolved on HMDA, 2,3-butanediol, adipate or isobutyrate were tolerant to other similar compounds not in the ALE set of compounds (mostly diamines, diols, diacids or monocarboxylic acids, respectively; Figure 3b). We found that in most cases strains tolerant to one compound also have improved growth rates on similar compounds, with an average growth improvement of 0.13 h^−1^ across the tested conditions (Figure 3b).

**Figure 3:**
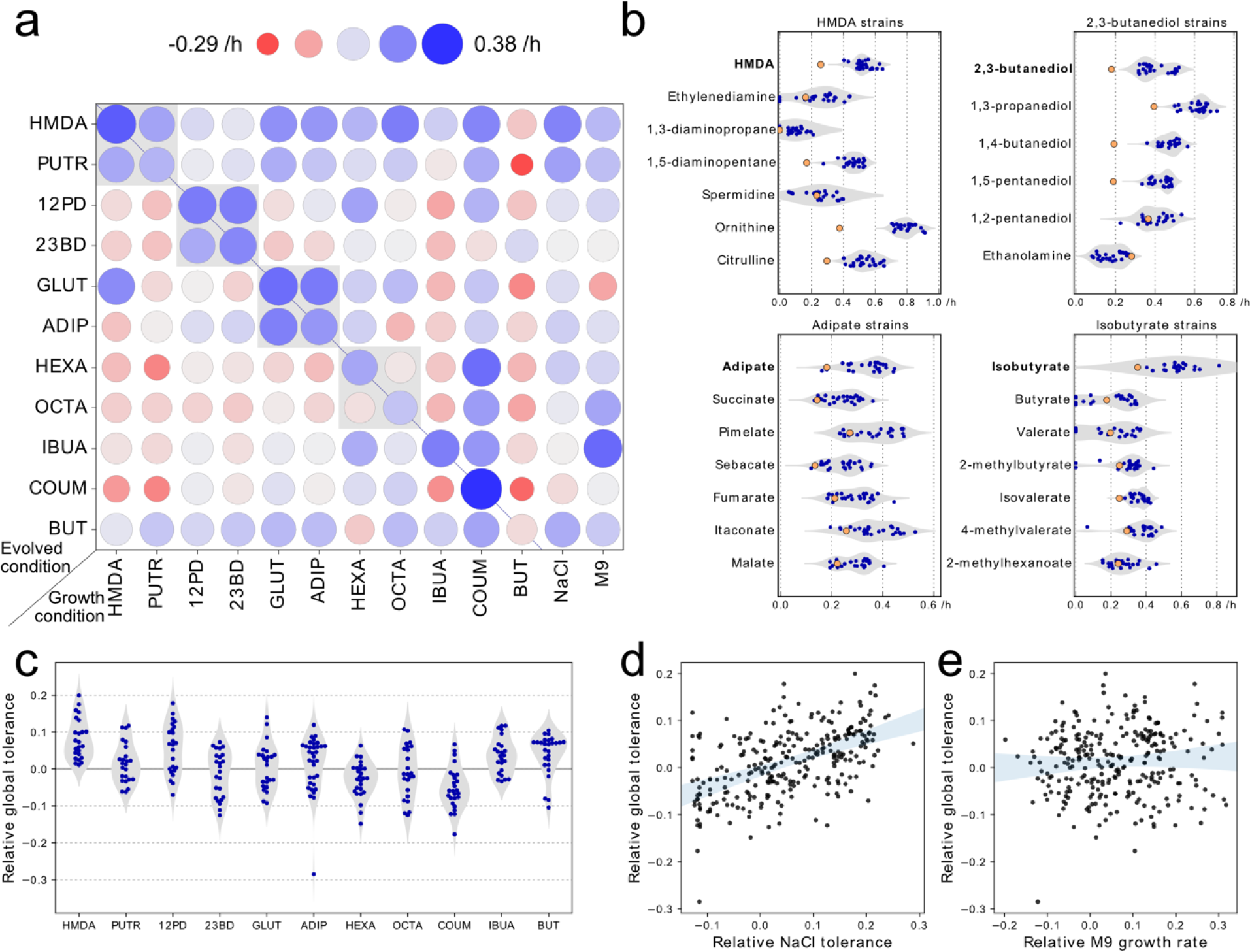
Cross-tolerance between similar and dissimilar compounds. a) Cross-tolerance between the compounds used for ALE. Circle color and size represent the mean growth rate of the group of strains relative to the unevolved reference strain. The grey boxes indicate pairs of compounds that are from similar chemical class. The growth rates on 0.6 M NaCl and M9 are also shown. b) Strains evolved in HMDA, 2,3-butanediol, adipate and isobutyrate conditions were screened for tolerance against other chemically similar compounds that were not part of the set of ALE compounds. Blue points represent growth rates of evolved strains, while the orange points show the growth rates of the reference strain. c) Distribution of global tolerance values (i.e. average relative growth rate across all 11 chemicals) for strains evolved on each of the 11 compounds d) Global tolerance as a function of osmotolerance (growth rate on NaCl) and e) as a function of improvement in baseline growth (growth on M9 glucose).

We sought to understand some of the general features that make *E. coli* tolerant to a broad range of chemicals. We used the average growth rate of an ALE strain relative to the wild-type strain across all 11 chemicals as a metric of global chemical tolerance of a strain. The global chemical tolerance of strains depended significantly on which chemical the specific strain had been evolved to tolerate (F = 10.06, p < 10^−13^; Figure 3c), and also varied between strains evolved to tolerate the same chemical with a median standard deviation of 0.06 h^−1^. Strains evolved on HMDA typically had high chemical tolerance whereas strains evolved on coumarate and hexanoate were less tolerant to most other chemicals than the wild-type strain. We found that NaCl tolerance was statistically significantly predictive of global chemical tolerance (Pearson’s r = 0.52, p < 10^−20^) (Figure 3d). In contrast, the relative growth rate of the ALE strain in M9 glucose was not predictive of global chemical tolerance (r = 0.06, p = 0.31) (Figure 3e) indicating that both general growth improvements and tolerance tradeoffs existed. The final osmolarity of the medium during ALE was not associated with either NaCl or global tolerance of the strain (Supplementary Figure 3).

### Tolerance mechanisms

For each chemical, we reintroduced combinations of some of the most commonly observed mutations into the background strain to determine causality. The mutations were introduced as gene deletions in cases where the observed mutations appeared likely to result in loss of function of the gene. We measured the tolerance of the resulting 145 genome edited strains with different combination of mutations to chemical stresses relevant for each strain (Figure 4a). Even though tolerance to most of the chemicals could be at least partially reconstructed by introducing up to four genome edits (Figure 4b), the mechanisms by which the mutations caused tolerance to each chemical were generally difficult to decipher from this data alone. Nonetheless, in some cases where we found mutations in the same gene in many independently evolved strains and these mutations conferred high level of tolerance in reconstructed strains, it was possible to formulate an experimentally testable mechanistic hypothesis.

All strains evolved to tolerate adipate and glutarate contained mutations in *kgtP*, which encodes an active alpha-ketoglutarate importer^14^. Approximately half of these mutations were clearly loss-of-function, i.e. frameshift mutations or single nucleotide polymorphisms (SNPs) that generated premature stop codons. We found that a *kgtP* deletion strain grew significantly faster than the wild-type strain in the presence of high levels of the diacids, particularly glutarate (Figure 4cd). Some of the diacid-evolved strains also contained loss-of-function mutations in two other transporter encoding genes, *proV* (subunit of the ProVWX glycine betaine transporter) and *ybjL* (uncharacterized putative transporter). Deleting these transporters (*proV* or *ybjL*) in addition to *kgtP* increased the growth rate further on glutarate and adipate (Figure 4c and 4d), with the triple deletion strain reaching the same growth rate on glutarate as the best evolved isolates.

**Figure 4:**
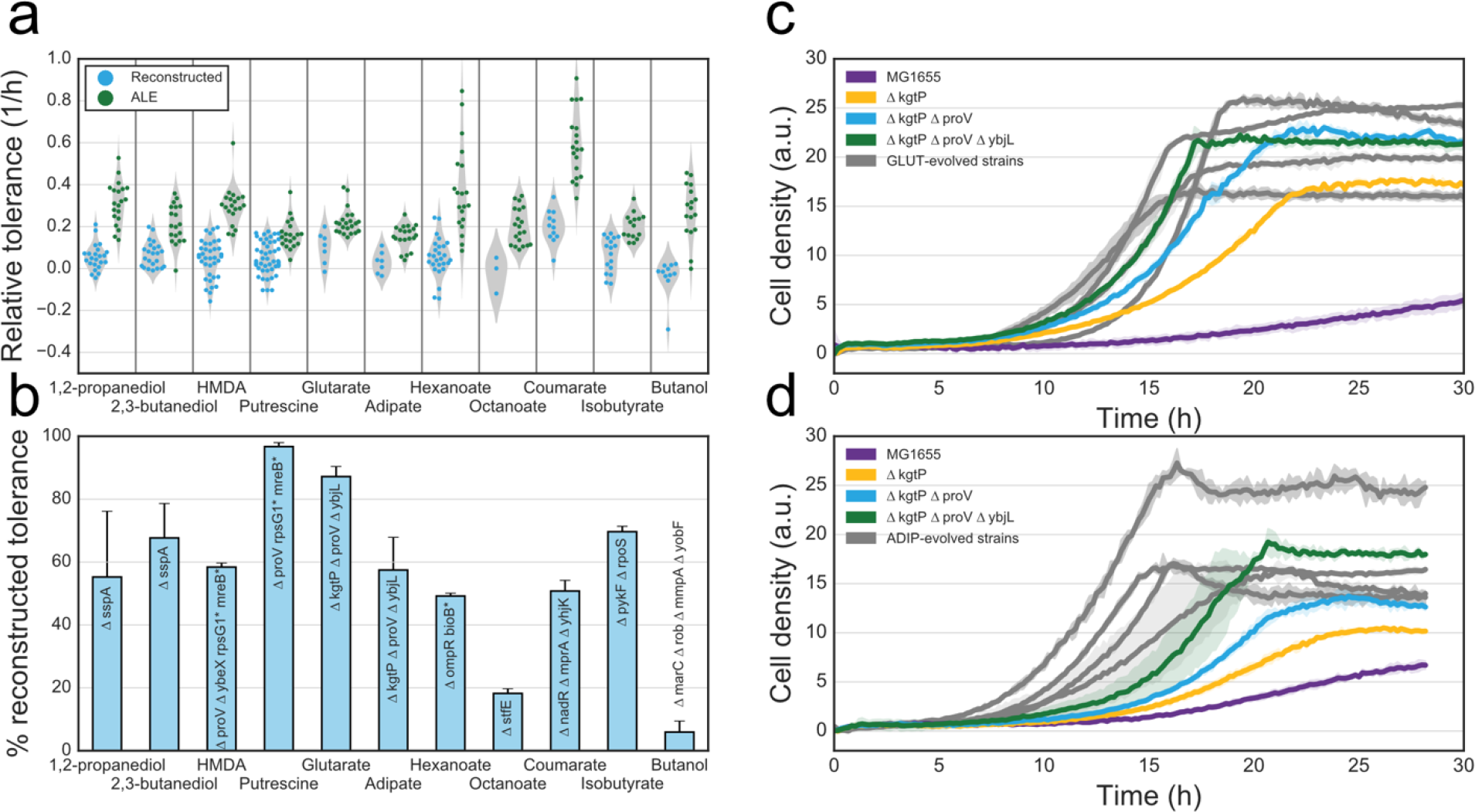
Elucidation of causative mutations. a) Distributions of tolerance to each of the 11 chemicals for reconstructed and evolved strains respectively. Relative tolerance was defined as the growth rate in presence of the given chemical relative to the growth rate of the reference strain. Tolerance for each specific reconstructed strain is shown in Supplementary Figure 6. b) The percentage of reconstructed tolerance to each of the chemicals, calculated as the relative tolerance of the best reconstructed strain divided by the 3^rd^ quartile of relative tolerances of the evolved strains. Error bars denote standard deviation of the growth rate of the the reconstructed strain. Asterisks denote that a point mutation has been inserted in the gene. See Supplementary Table 3 for a list of reconstructed strain genotypes. c) Growth curves of reference strain MG1655, four genetically distinct glutarate-evolved strains as well as transporter deletion strains in M9 glucose with 47.5 g/L glutarate. d) Growth curves of the reference strain MG1655, four genetically distinct adipate-evolved strains as well as transporter deletion strains in M9 glucose with 50 g/L adipate. Lines show mean growth curves of 3 biological replicates, with error bars indicating the standard deviation about the mean.

### Production in evolved strains

To determine whether strains evolved to tolerate a non-native product would produce more of the corresponding product, we inserted production pathways into the set of ALE-derived trains. We chose the two pyruvate-derived compounds, isobutyrate and 2,3-butanediol, as examples because the two tolerized sets of strains had very different genotypes and growth phenotypes from each other (Table 1 and Figure 3), and production of these compounds has previously been demonstrated in *E. coli*^15,16^.

We introduced an isobutyrate production pathway^15^ into wild-type MG1655 and 12 genetically distinct isobutyrate-tolerant strains by expressing three heterologous genes from plasmids and deleting a competing pathway in each strain (Figure 5a). The engineered ALE-derived strains had highly variable levels of production of isobutyrate (Figure 5b), with some strains producing almost no isobutyrate and the best ALE-derived strain producing over three times more isobutyrate than the engineered wild-type strain. The best producers (IBUA8-3 and IBUA8-10) both had *ilvH/N* mutations, which likely reduce valine feedback inhibition of the first acetolactate synthase step of branched chain amino acid biosynthesis^17^ (Supplementary Figure 4).

We also introduced a 2,3-butanediol production pathway^16^ into MG1655 and 20 ALE-derived strains by expressing three heterologous genes in the strains (Figure 5c). Again, there was variation in 2,3-butanediol production among the engineered ALE strains, but the majority of strains had production levels similar to the engineered wild-type strain and only two ALE strains showed a significant improvement in production of 2,3-butanediol compared to the engineered reference strain (Figure 5d). Although comparison to other isolates from the same populations allowed us to identify the mutations responsible for improved production, we could not identify a mechanistic basis for this (see Supplementary results and discussion).

**Figure 5:**
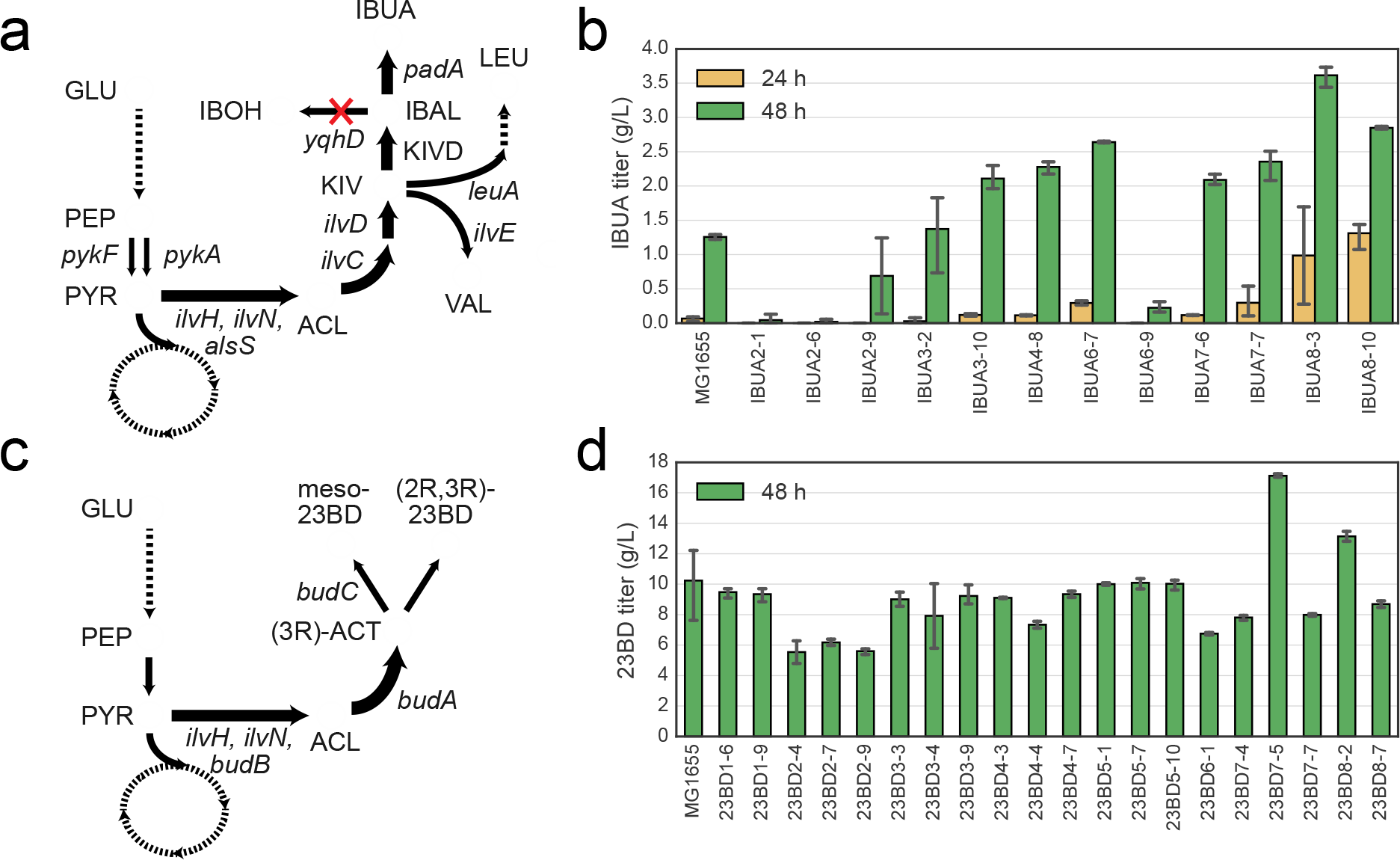
Production of isobutyric acid and 2,3-butanediol using ALE-derived tolerized strains. a) The production pathway schematic for isobutyrate, with heterologous expression of an acetolactate synthase AlsS, ketoisovalerate decarboxylase KIVD, and PadA to generate isobutyric acid from ketoisovalerate (KIV), with deletion of native yqhD to prevent reduction of isobutyraldehyde (IBAL) to isobutanol (IBOH). b) Production of isobutyrate in wild-type and evolved isolates harboring production plasmids for isobutyrate and deletion of yqhD after 24 and 48 hours growth in FIT (feed-in-time) medium. c) Production pathway schematic for 2,3-butanediol from pyruvate, with heterologous expression of BudA, BudB, and BudC. d) Production of 2,3-butanediol in wild-type and evolved isolates harboring a production plasmid for 2,3-butanediol and deletion of hsdR after 48 hours in M9 + 5% glucose + 0.5% yeast extract.

## Discussion

Our results demonstrated that ALE can be used to increase the tolerance of microbial cells to an exogenously supplied chemical. The tolerated concentrations increased 60-400%, with the largest increases seen for chemicals that initially were most toxic to *E. coli*, while tolerance to compounds that were initially tolerated at high levels, such as diols, increased more modestly. In comparison to previous ALE studies of chemical tolerance^5^, the systematic approach used here enables direct comparisons of the evolvability of *E. coli* tolerance towards different chemical stresses. A similar automated ALE approach has been previously taken to study adaptation to diverse stresses including some chemical stresses in *E. coli*^18^, but in the present study we used significantly higher concentrations of chemicals to mimic industrially relevant conditions and evolved more populations per condition.

Resequencing of 224 evolved strains revealed relatively low numbers of mutations in most strains. In principle, this facilitates a good understanding of which genes are important for tolerance to a given compound. However, the observed genomic variant landscape across all strains was complex, with most mutations found only in specific evolution conditions, thereby indicating a high diversity in mechanisms of toxicity and tolerance between the compounds. Interpretation of the resequencing data was also complicated by the high frequency of mutations in regulatory genes with known pleiotropic effects. The overall conclusions from the resequencing results is that there are no universal tolerance mechanisms at the genetic level for this set of chemicals and that tolerance usually involved both specific (e.g. transporters) and general (e.g. adjustments in global regulation) adaptations (Supplementary Results and Discussion).

Cross-tolerance profiling showed that strains evolved to tolerate one diacid, diol, or diamine also had tolerance to the other chosen chemical of the same class (Figure 3a). Furthermore, strains evolved to tolerate a specific chemical tended to be tolerant to a wide range of similar chemicals (Figure 3b). This is of great practical relevance, as it is potentially only necessary to perform ALE once for a class of chemicals (e.g. diols) in order to obtain a series of platform strains that have high levels of tolerance within the chemical class.

Cross tolerance profiling could be used to define a measure of global chemical tolerance of each evolved strain, which was found to be highly variable between genetically distinct strains evolved in one condition and even more so between strains evolved in different conditions. We found global tolerance to be uncorrelated to the growth rate in M9 glucose medium (Figure 3d) indicating that fast growth and stress tolerance are not always correlated. Although previous studies have found that evolving *E. coli* to in M9 medium collaterally reduced stress tolerance^19^, this tradeoff does not seem to apply in the reverse direction for the set of chemicals used here, as most evolved tolerant strains grew faster in M9 than the wild-type strain. On the other hand, the ability of a strain to grow in NaCl was found to be significantly predictive of the global chemical tolerance of the strain (Figure 3d), suggesting that non-specific osmotolerance (or cation tolerance to Na+^20^) explains part of the observed increases in tolerance. Since acids and diacids were neutralized with NaOH in this study, tolerance to Na+ is likely driving part of the correlation seen in Figure 3d. Some highly tolerant strains had mutations in genes such as *nagC* and *proV* that have previously been implicated in NaCl tolerance in ALE experiments^13^. Unfortunately, the exact mechanisms by which many of the observed mutations confer NaCl tolerance remain elusive. Several of the most broadly tolerant strains were from the HMDA condition (Figure 3), however these strains had not evolved significant tolerance towards the diols, even though those conditions also had very high osmolarity. This dissociation between tolerance to cation stress and non-ionic osmotic stress has also been observed previously^20^.

Determining the exact mechanisms of chemical tolerance was challenging, but in specific cases convergent mutation targets allowed mechanistic hypotheses to be generated and tested. Since all strains evolved on adipate and glutarate contained mutations in the *kgtP* gene encoding an alpha-ketoglutarate transporter and given the structural similarity of glutarate and adipate to alpha-ketoglutarate, this was likely the primary importer of the two diacids. Indeed, deletion of *kgtP* conferred a large increase in diacid tolerance. Two further transporters, *proV* and *ybjL*, were mutated in specific diacid-tolerant strains and a triple deletion of these transporters was sufficient to achieve levels of tolerance to glutarate and adipate on par with the evolved strains. The *proV* gene encodes a subunit of the ProVWX complex that imports the osmoprotectant glycine betaine^21^. As ProVWX mutations were widely observed, it is possible that this transporter, which is highly expressed in osmotic stress conditions, simply imposes a burden on the cell in these conditions^22^.

To investigate if pre-evolving for exogenous tolerance could improve endogenous production, pathways for isobutyrate and 2,3-butanediol were engineered into strains that had been evolved to tolerate the respective compounds. The engineered ALE strains did not generally show increased production, but for both compounds, we could identify specific strains that had significantly higher production than the corresponding engineered wild-type strain. This indicated that evolving for exogenous tolerance could be a viable strategy for obtaining improved production strains as long as a sufficient number of independently evolved strains are engineered and screened for production.

Studying the ALE-derived strains that had increased production of isobutyrate or 2,3-butanediol allowed us to identify roles for specific mutations in enhancing production. All isobutyrate-evolved strains contained *pykF* mutations, the majority of which were clear loss-of-function mutations. Mutations in *pykF* are commonly seen in many *E. coli* ALE experiments^12,23^ and *pykF* deletion has also been shown to allow increased production of many metabolites^24,25^. Deletion of *pykF* has been shown to redirect fluxes in central carbon metabolism and result in reduced intracellular pyruvate levels^26^. While *pykF* deletion significantly improved tolerance to isobutyrate (Supplementary Figure 4), the different ALE strains showed widely varying production capabilities ranging from no production to three times higher than the reference strain, indicating that *pykF* mutations alone did not strongly affect production. The highest producing ALE-derived strains had mutations in *ilvH/N* encoding regulatory acetolactate synthase (ALS) subunits. These mutations were shown to alleviate feedback inhibition by valine (Supplementary Figure 4) that results in isoleucine starvation^27^ if valine levels increase. This may explain their ability to produce higher levels of isobutyrate, as the engineered strains contain a heterologously expressed ALS which may further increase valine levels relative to isoleucine. In the case of 2,3-butanediol production, only one of the engineered evolved strains (23BD7-5) had considerably higher production than the wild-type strain. Compared with lower production strains from the same population, this strain had loss-of-function mutations in the *acrB* (encoding a subunit of the AcrAB-TolC multidrug efflux pump) and *purT* (encoding a phosphoribosylglycinamide formyltransferase) genes. Deletion of *acrA* or *acrB* has previously been shown to increase tolerance towards isobutanol^10^, but it is not clear why either *acrB* or *purT* mutations should increase 2,3-butanediol production.

In conclusion, the results of this study showed that *E. coli* can be evolved to tolerate high concentrations of a wide range of industrially relevant chemicals. A strain that is evolved to tolerate one chemical is likely to also have increased tolerance to other chemically similar compounds, allowing evolved strains to be used as platform strains for production of several different chemicals. Strains that are tolerant to NaCl tend to be tolerant to most chemicals at high concentrations, but strains that growth rapidly on M9 glucose minimal media do not necessarily exhibit broad chemical tolerance. Additionally, we have shown that evolving strains to tolerate a chemical can have beneficial effects on the strain’s ability to produce the chemical, further demonstrating the value of ALE during strains development for microbial production.

## Supporting information

Methods and Supplementary Results and Discussion

## Acknowledgements

The authors acknowledge funding from the Novo Nordisk Foundation (Grant numbers NNF10CC1016517 and NNF140C0011269). The authors thank Katy Kao and Hannes Link for fruitful discussions.

## References

1. Van Dien, S. From the first drop to the first truckload: Commercialization of microbial processes for renewable chemicals. Curr. Opin. Biotechnol. 24, 1061–1068 (2013).

2. Deparis, Q., Claes, A., Foulquié-Moreno, M. R. & Thevelein, J. M. Engineering tolerance to industrially relevant stress factors in yeast cell factories. FEMS Yeast Res. 17, 1–17 (2017).

3. Qi, Y., Liu, H., Chen, X. & Liu, L. Engineering microbial membranes to increase stress tolerance of industrial strains. Metab. Eng. 53, 24–34 (2019).

4. Hansen, A. S. L., Lennen, R. M., Sonnenschein, N. & Herrgård, M. J. Systems biology solutions for biochemical production challenges. Curr. Opin. Biotechnol. 45, 85–91 (2017).

5. Winkler, J. D. & Kao, K. C. Recent advances in the evolutionary engineering of industrial biocatalysts. Genomics 104, 406–411 (2014).

6. Kildegaard, K. R. et al. Evolution reveals a glutathione-dependent mechanism of 3-hydroxypropionic acid tolerance. Metab. Eng. 26, 57–66 (2014).

7. Haft, R. J. F. et al. Correcting direct effects of ethanol on translation and transcription machinery confers ethanol tolerance in bacteria. Proc. Natl. Acad. Sci. 111, E2576--E2585 (2014).

8. Reyes, L. H., Abdelaal, A. S. & Kao, K. C. Genetic determinants for n-butanol tolerance in evolved escherichia coli mutants: Cross adaptation and antagonistic pleiotropy between n-butanol and other stressors. Appl. Environ. Microbiol. 79, 5313–5320 (2013).

9. Mundhada, H. et al. Increased production of L-serine in Escherichia coli through Adaptive Laboratory Evolution. Metab. Eng. 39, 141–150 (2017).

10. Atsumi, S. et al. Evolution, genomic analysis, and reconstruction of isobutanol tolerance in Escherichia coli. Mol. Syst. Biol. 6, 1–11 (2010).

11. Royce, L. A. et al. Evolution for exogenous octanoic acid tolerance improves carboxylic acid production and membrane integrity. Metab. Eng. 29, 180–188 (2015).

12. Wang, X., Zorraquino, V., Kim, M., Tsoukalas, A. & Tagkopoulos, I. Predicting the evolution of Escherichia coli by a data-driven approach. Nat. Commun. 9, (2018).

13. Winkler, J. D., Garcia, C., Olson, M., Callaway, E. & Kao, K. C. Evolved osmotolerant escherichia coli mutants frequently exhibit defective N-acetylglucosamine catabolism and point mutations in cell shape-regulating protein MreB. Appl. Environ. Microbiol. 80, 3729–3740 (2014).

14. Seol, W. & Shatkin, A. J. Escherichia coli kgtP encodes an alpha-ketoglutarate transporter. Proc. Natl. Acad. Sci. 88, 3802–3806 (1991).

15. Zhang, K., Woodruff, A. P., Xiong, M., Zhou, J. & Dhande, Y. K. A synthetic metabolic pathway for production of the platform chemical isobutyric acid. ChemSusChem 4, 1068–1070 (2011).

16. Xu, Y. et al. Systematic metabolic engineering of Escherichia coli for high-yield production of fuel bio-chemical 2,3-butanediol. Metab. Eng. 23, 22–33 (2014).

17. Kaplun, A. et al. Structure of the Regulatory Subunit of Acetohydroxyacid Synthase Isozyme III from Escherichia coli. 951–963 (2006). doi:10.1016/j.jmb.2005.12.077

18. Horinouchi, T. et al. Prediction of Cross-resistance and Collateral Sensitivity by Gene Expression profiles and Genomic Mutations. Sci. Rep. 7, 1–11 (2017).

19. Utrilla, J. et al. Global Rebalancing of Cellular Resources by Pleiotropic Point Mutations Illustrates a Multi-scale Mechanism of Adaptive Evolution. Cell Syst. 2, 260–271 (2016).

20. Wu, X., Altman, R., Eiteman, M. A. & Altman, E. Adaptation of Escherichia coli to Elevated Sodium Concentrations Increases Cation Tolerance and Enables Greater Lactic Acid Production. Appl. Environ. Microbiol. 80, 2880–2888 (2014).

21. Wood, J. M. Bacterial Osmosensing Transporters. Methods in Enzymology: Osmosensing and Osmosignaling 428, (Elsevier Masson SAS, 2007).

22. Gunasekera, T. S., Csonka, L. N. & Paliy, O. Genome-wide transcriptional responses of Escherichia coli K-12 to continuous osmotic and heat stresses. J. Bacteriol. 190, 3712–3720 (2008).

23. Phaneuf, P. V, Gosting, D., Palsson, B. O. & Feist, A. M. ALEdb 1.0: A Database of Mutations from Adaptive Laboratory Evolution Experimentation. Nucleic Acids Res. 1–8 (2018). doi:10.1093/nar/gky983

24. Sengupta, S., Jonnalagadda, S., Goonewardena, L. & Juturu, V. Metabolic engineering of a novel muconic acid biosynthesis pathway via 4-hydroxybenzoic acid in Escherichia coli. Appl. Environ. Microbiol. 81, 8037–8043 (2015).

25. Harder, B. J., Bettenbrock, K. & Klamt, S. Model-based metabolic engineering enables high yield itaconic acid production by Escherichia coli. Metab. Eng. 38, 29–37 (2016).

26. Al Zaid Siddiquee, K., Arauzo-Bravo, M. J. & Shimizu, K. Metabolic flux analysis of pykF gene knockout Escherichia coli based on 13C-labeling experiments together with measurements of enzyme activities and intracellular metabolite concentrations. Appl. Microbiol. Biotechnol. 63, 407–417 (2004).

27. De Felice, M., Levinthal, M., Iaccarino, M. & Guardiola, J. Growth inhibition as a consequence of antagonism between related amino acids: Effect of valine in Escherichia coli K-12. Microbiol. Rev. 43, 42–58 (1979).

