## Supplementary material for "Adaptive laboratory evolution reveals general and specific chemical tolerance mechanisms and enhances biochemical production": Methods and Supplementary Results and Discussion

### Online supplementary material

#### Methods

##### Strains and media

*E. coli* K-12 MG1655 strain was used as a starting point strain for the adaptive laboratory evolution experiments and as reference strain for all subsequent characterization. Mutations present in the parental strain compared with the public reference sequence can be found at https://aledb.org/mutations/?ale_experiment_id=116. Chemicals were purchased from Sigma-Aldrich (Merck KGaA, Darmstadt, Germany), Fisher Scientific (Part of Thermo-Fisher Scientific) or TCI (TCI Europe N.V., Zwijndrecht, Belgium). Plasmids for isobutyric acid and 2,3-butanediol production were obtained from the authors [1,2].

M9 glucose medium supplemented with 10 g/L glucose was formulated with 1x M9 salts, 2 mM $MgSO_4$, 100 µM $CaCl_2$ and 1x trace elements. A stock solution of 10x M9 salts consisted of 68 g/L $Na_2HPO_4$ anhydrous, 30 g/L $KH_2PO_4$, 5 g/L NaCl, and 10 g/L $NH_4Cl$ dissolved in Milli-Q filtered water and autoclaved. M9 trace elements stock concentration was a 2000x solution containing of 3.0 g/L $FeSO_4 \cdot 7H_2O$, 4.5 g/L $ZnSO_4 \cdot 7H_2O$, 0.3 g/L $CoCl_2 \cdot 6H_2O$, 0.4 g/L $Na_2MoO_4 \cdot 2H_2O$, 4.5 g/L $CaCl_2 \cdot H_2O$, 0.2 g/L $CuSO_4 \cdot 2H_2O$, 1.0 g/L $H_3BO_3$, 15 g/L disodium EDTA, 0.1 g/L KI, 0.7 g/L $MnCl_2 \cdot 4H_2O$ in Milli-Q filtered water and sterile filtered.

All laboratory evolutions, screening and cultivation experiments were done at neutral pH. Acids that were not sodium salts were neutralized with NaOH. They were brought to pH 7.0 with base as more concentrated stocks in water and diluted in the final M9 media stocks to the correct concentration as indicated in Supplementary Table 1.

##### Selection of initial chemical concentrations

The toxicity of each of the chemicals was tested by screening growth of MG1655 in different concentrations. Biological triplicates of *E. coli* MG1655 were cultivated at 37 °C with 300 rpm shaking. After 14 to 18 h, the cultures were inoculated into M9 + 0.2 % glucose and one of the chosen chemicals at different concentrations at neutral pH. The cultures were then incubated in a BioLector microbioreactor system (m2p-labs GmbH, Baesweiler, Germany) at 37 °C with 1000 rpm shaking. The growth rates at different concentrations were calculated for each chemical (Supplementary Figure 1). Initial concentrations for adaptive laboratory evolution were chosen so that MG1655 could obtain a growth rate of approximately 0.4 $h^{-1}$.

The initial and final evolution concentrations, as well as the screening concentrations for all the chosen chemicals are shown in Supplementary Table 1.

#### Adaptive laboratory evolution

The starting strain K-12 MG1655 was adaptively evolved for higher concentrations of each chemical through independent parallel replicates. Bacterial cells were cultivated in M9 + glucose supplemented with the initial chemical concentration listed in Supplementary Table 1, with gradual increase in each chemical concentration over the time span of the laboratory evolution experiment. The growth medium was kept at neutral pH to ensure an evolutionary pressure for tolerating the specific chemical compounds rather than tolerance towards low or high pH. Cells were serially passaged during exponential growth for approximately 40 days using an automated liquid-handler platform as described by LaCroix et al. (2017). Cells were cultured at 37 °C with full aeration at 1200 rpm stirring speed. Once $OD_{600}$ reached approximately 1.0, 150 µL was passed into a new tube with 15 mL fresh media containing the respective chemical concentration. Over the course of the experiment, cells were kept in exponential growth phase in order to keep a constant selection pressure for growth rate. The $OD_{600}$ was measured by a Sunrise Plate Reader (Tecan Inc., Switzerland). Growth rates were determined by computing the slope of log(OD) using linear regression with the Polyfit function in MATLAB (The Mathworks Inc., Natick, Massachusetts). When an increase in the apparent growth rate was achieved (average growth rate for all of the parallel replicates) at a particular concentration, the chemical concentration was increased by 10-15%. This process was repeated in cycles until a significant increase in tolerated concentration was achieved. In incidents where the increase in the chemical concentration caused cells to crash, i.e. cell death, chemical concentration was reduced to a level that allowed cell growth. Periodically, samples were frozen in 25% v/v glycerol and stored at -80 °C for further use.

#### Growth screening of ALE isolates

##### Primary tolerance screening

Populations from evolution endpoints were plated on LB agar plates (or on M9 agar in the rare cases where growth was not observed on LB agar) and 10 individual colonies from each population were screened for growth at the maximum concentration for which robust growth rates were achieved during the evolution (Supplementary Table 1). Cultures of the wild-type strain, *E. coli* K-12 MG1655, were used as controls. The isolates were inoculated in 300 µL M9 + glucose in deep-well plates and incubated in plate shaker at 37 °C and 300 rpm. The next day, cells were diluted 10X in M9 + glucose and 30 µL was transferred to clear-bottom 96 half-deep plates containing 270 µL M9 + glucose supplemented with the corresponding toxic chemical at concentrations as in Supplementary Table 1 at neutral pH (isolate screening concentration). The plates were incubated at 37 °C with 225 RPM shaking in a Growth Profiler screening platform (EnzyScreen BV, Heemstede, Netherlands). The resulting growth curves for all 880 isolates were inspected qualitatively for isolates exhibiting robust growth as assessed by lag time, final OD and growth rate. Each of the 10 isolates per population from the primary screening were grouped according to close similarities based on the above criteria. For each population, isolates representative of each group were picked (2-3 isolates per population).

##### Cross-tolerance screening

The *E. coli* strains were inoculated into 300 µl of M9 + glucose medium in 96 deep well plates (in biological triplicates, except for 5 biological replicates for K-12 MG1655) and the cultures were incubated at 37°C and 300 rpm overnight. The next day, 30 µl of a 10-fold dilution in M9 + glucose was added to 270 µl of M9 + glucose supplemented with chemicals at neutral pH in a 96 well plate format, and the plates were incubated in growth profiler (EnzyScreen BV, Heemstede, The Netherlands) at 37 °C and 225 RPM. The chemical concentrations are shown in Supplementary Table 1 ("Cross-tolerance screening concentration"). For cross-tolerance screening the chemical concentrations were lower than those used for primary ALE isolate screening above in order to allow for the wild type control strain to grow in all conditions.

#### Genome editing of *E.coli*

Strains containing the relevant single gene deletions were obtained from the Keio Collection and were transduced into the MG1655 background strain using a protocol previously described[4]. Multiple gene deletions were constructed using a protocol previously described[4]. Site directed changes in the *E. coli* genome of evolved strains were made using a previously described protocol[5].

#### 2,3-butanediol production

The *hsdR* gene was deleted from each of the strains evolved on 2,3-butanediol, since it was found that the production pathway plasmid (pET-RABC) contained an EcoK restriction site. The strains were then transformed with pET-RABC[2] and precultured in 300 µl of M9 + glucose supplemented with 5 g/L yeast extract and kanamycin (50 µg/mL) in a 96 deep well plate and incubated at 37 °C with 300 RPM overnight (incubated for 20 h) in quadruplicates. *E. coli* K-12 MG1655 Δ*hsdR*/pET-RABC was used as a control. The following day, 20 µL of pre-inoculum was transferred into 2 mL of the same medium in 24 deep well plates and incubated at 30 °C and 300 RPM. At 24 h and 48 h, optical densities of the culture broths were determined at 600 nm ($OD_{600nm}$). Then, 400 µL of the cultures were harvested, centrifuged at 4000 RPM for 10 min and 30 µL of the collected supernatants were injected into high performance liquid chromatography (HPLC).

The amounts of 2,3-butanediol in the supernatants were quantified by HPLC (Ultimate 3000, Thermo Scientific, USA) equipped with an organic acid analysis column, Aminex® HPX-87H ion exclusion column (300 mm x 7.8 mm, Bio-Rad Laboratories, Denmark) connected to a refractive Index (RI) detector and a UV detector (205 nm, 210 nm, 254 nm and 280 nm). An isocratic elution with flow rate of 0.5 mL/min of 5 mM sulphuric acid was used for 30 min. Under these conditions, stereoisomers of 2,3-butanediol were detected under the RI detector channel at the retention times of 17.4 min and 18.3 min. Using the peak areas of the stereoisomers, total amount of 2,3-butanediol was calculated. For absolute quantification a calibration curve was drawn using 1, 5, 10, 12.5, 15 and 25 g/L concentrations ($y = 6.5119x + 0.5464$, $R^2 = 0.9999$).

#### Isobutyrate production

The *yqhD* gene was deleted from each of the strains evolved on isobutyrate. The strains were then transformed with pIBA1 and pIBA7 plasmids[1] and precultured into 300 µl of LB media supplemented with kanamycin (50 µg/mL) and ampicillin (100 µg/mL) in 96 deep well plate and incubated at 37 ºC with 300 RPM overnight (incubated for 18 h) (in quadruplicates). *E. coli* MG1655 Δ*yqhD*/pIBA1/pIBA7 was used as a control. The following day, 24 µL of pre-inoculum was transferred into 2.4 mL of half-FIT media (1:1 FIT media: 200 mM MOPS) supplemented with antibiotics. Then the culture plates were incubated at 30 °C and 300 RPM. After 6 hours of incubation, $OD_{600}$ was measured and the cultures were induced with 100 µM of IPTG and continued the incubation at 30 °C and 300 RPM. At 24 h, 48 and 72 h, $OD_{600}$ was measured again. Then, 300 µl of the cultures were harvested, centrifuged at 4000 RPM for 10 min and 30 µL of the collected supernatants were injected into HPLC.

The amounts of isobutyrate in the supernatants were quantified by HPLC (Ultimate 3000, Thermo Scientific, USA) equipped with an organic acid analysis column, Aminex® HPX-87H ion exclusion column (300 mm x 7.8 mm, Bio-Rad Laboratories, Denmark) connected to a refractive Index (RI) detector and a UV detector (205 nm, 210 nm, 254 nm and 280 nm). An isocratic elution with flow rate of 0.5 mL $min^{-1}$ of 5 mM sulphuric acid was used for 30 min. Under these conditions, isobutyrate was detected under the 210 nm UV channel at a retention time of 20.3 min. Using the peak area, the total amount of isobutyrate was calculated. For absolute quantification a calibration curve was drawn using 0.5, 1, 2.5, 4, 5, 7.5 10, and 12.5 g/L concentrations ($y = 35.487x - 2.3142$, $R^2 = 0.9993$).

#### Resequencing

Genomic libraries were generated using the TruSeq® Nano DNA LT Library Prep Kit (Illumina Inc., San Diego CA). Briefly, 100 ng of genomic DNA diluted in 52.5 µL TE buffer was fragmented in Covaris Crimp Cap microtubes on a Covaris E220 ultrasonicator (Woburn, MA) with 5% duty factor, 175 W peak incident power, 200 cycles/burst, and 50-s duration under frequency sweeping mode at 5.5 to 6°C (Illumina recommendations for a 350-bp average fragment size). The ends of fragmented DNA were repaired by T4 DNA polymerase, Klenow DNA polymerase, and T4 polynucleotide kinase. The Klenow exo minus enzyme was then used to add an 'A' base to the 3' end of the DNA fragments.

The adapters were ligated to the ends of the DNA fragments, and the DNA fragments ranging from 300 - 400 bp were recovered by beads purification. Finally, the adapter-modified DNA fragments were enriched by 3 cycle PCR. Final concentration of each library was measured by Qubit® 2.0 Fluorimeter and Qubit DNA Broad range assay (Life Technologies). Average dsDNA library size was determined using the Agilent DNA 7500 kit on an Agilent 2100 Bioanalyzer. Libraries were normalized and pooled in 10 mM Tris-Cl, pH 8.0, plus 0.05% Tween 20 to the final concentration of 10 nM. Denaturated in 0.2N NaOH, 10 pm pool of 20 libraries in 600 µL ice-cold HT1 buffer was loaded onto the flow cell provided in the MiSeq Reagent kit v2 (300 cycles) (Illumina Inc., San Diego CA) 300 cycles and sequenced on a MiSeq (Illumina Inc., San Diego CA) platform with a paired-end protocol and read lengths of 151 nt.

#### Resequencing data analysis

The Illumina sequencing reads were analyzed with the Breseq pipeline[6] through the ALEdb platform[7] to generate lists of mutations for each evolved strain. The reference strain for this analysis was *E. coli* K-12 MG1655 with the Genbank accession number NC_000913.3. The variant calling data is available through the public ALEdb platform at http://aledb.org.

Each mutation was mapped to one or more genes. Intragenic mutations were mapped to any gene(s) whose coding sequence overlapped with the mutation. Intergenic mutations were mapped to the closest gene downstream from the mutation.

#### Growth data analysis

The growth curves generated by the instruments were processed using the *croissance* python package (http://github.com/biosustain/croissance), which performs automated growth phase identification and growth parameter fitting. Biomass concentration was quantified by $OD_{600}$ values. For each extracted growth rate, a normalized growth rate was calculated by subtracting the mean growth rate of the wild-type strain, MG1655, on the same plate and in the same medium. This was done to remove the effects of any between-plate and between-experiment growth variations.

The *croissance* algorithm consists of two separate steps:

**Step 1:** The growth curve is smoothed and analyzed to find regions of exponential growth. This is done by identifying time intervals where the first- and second-order time derivatives of the smoothed biomass function are strictly positive.

**Step 2:** Each growth phase identified in step 1 is fitted with an exponential function of the form

$$x(t) = a \cdot e^{\mu \cdot t} + b$$

where $\mu$ (growth rate) is of particular interest in this study. The offset parameter $b$ is included to enable analysis of growth curves that are not background-subtracted.

Post-processing was done to filter the returned growth rates. This served both to exclude growth rates from growth phases that were not thought to be real, and to select between several growth phases in the same growth curve. Growth rates higher than 1.5 $h^{-1}$ were excluded, as were growth phases where the absolute value of the fitted offset parameter *b* was larger than a certain threshold, *c* (0.5 for growth profiler curves, 4 for Biolector curves). Growth phases where the initial biomass concentration deviated more than *c* from the fitted offset *b* were also excluded, as these were likely to be secondary growth phases. Furthermore, growth curves starting after a certain time point were also excluded. This was done to prevent growth from contaminations from being used in the analyses. The chosen time cutoff was dependent on the growth conditions and varied between 30 and 40 hours. Very short growth phases were also excluded as they were most likely artifacts. For standard M9 glucose cultures growth phases shorter than 2 hours were excluded, while the cutoff was 5 hours for cultures in the stress conditions.

### Supplementary Results and Discussion

Below we briefly discuss some of the major mechanisms of tolerance that could be inferred from the resequencing data (http://aledb.org and Supplementary Tables 2,4 and 6) and growth phenotyping of strain reconstructions (Supplementary Table 3 and Supplementary Figure 6). We focus on genetic mechanisms that are shared by the majority of evolved populations or mechanisms that the strain reconstructions showed to contribute significantly to tolerance. In many of the conditions, there are diverse mechanisms by which tolerance is achieved and our reconstructions did not attempt to study all of the mechanisms. It was often not possible determine the mechanisms of chemical toxicity from the data collected in this study, as most of the mutations did not occur in direct molecular targets of chemical toxicity.

#### Evolved tolerance against diacids

As discussed in the main text, tolerance towards diacids (glutarate and adipate) was highly associated with mutated transporters. The *kgtP* gene was mutated in all isolates from either of the two conditions, and a *kgtP* knockout single knockout strain had increased tolerance to both glutarate and adipate (Supplementary Figure 6). Given the *kgtP* product's known function as an alpha-ketoglutarate importer, it is likely that the diacids are promiscuously imported by the transporter. A high frequency of mutations was also observed in the transporter-encoding genes *proV* and *ybjL*, and deletion of these transporters, in addition to *kgtP,* conferred higher levels of tolerance to both glutarate and adipate (Supplementary Figure 6). We found that the strains in which *kgtP* was mutated or deleted did not grow with glutarate as a carbon source whereas the wild-type and *proV* and *ybjL* single deletion strains did grow in with glutarate as a sole carbon source (data not shown), supporting the hypothesis that *kgtP* can promiscuously transport glutarate into the cell.

#### Evolved tolerance against isobutyrate

Isobutyrate-evolved strains were discussed in the main text and we provide some additional details on the tolerance mechanisms here. Most isolates from the isobutyrate condition had apparent loss-of-function mutations in the pyruvate kinase-encoding *pykF* gene, suggesting it is key in the evolution of tolerance towards isobutyrate. Deletion of *pykF* alone was also sufficient to allow quite a high level of tolerance to isobutyrate (Supplementary Figure 4A). It is not clear why *pykF* would be beneficial for isobutyrate tolerance as the effects of *pykF* deletion are quite broad including increasing glucose uptake rate and increasing NADPH production through the pentose phosphate pathway.

As explained in the main text *pykF* loss-of-function mutations did not necessarily allow increased production of isobutyrate – this required the presence of additional mutations in the *ilvH/N* regulatory subunits of the acetolactate synthase enzyme (ALS) that controls entry into branched-chain (and isobutyrate) biosynthetic pathways. We tested the effect of the ilvH mutation (L9F) in isolation and found that it did not alone significantly improve isobutyrate tolerance (Supplementary Figure 4A). On the other hand, our data showed *ilvH (L9F)* mutation alone was able to alleviate valine inhibition of growth (Supplementary Figure 4B) and this inhibition was also alleviated in a isobutyrate-evolved isolate (IBUA8-3) which had this mutation. Our data does not allow us to conclude what the mechanism of isobutyrate toxicity is, but one possibility is that isobutyrate inhibits a biosynthetic step downstream of ketoisovaleric acid (KIV in Figure 5) resulting in overflow of valine, which in turn results in inhibition of ALS. This finally causes a reduction in isoleucine supply and slow growth phenotype.

#### Evolved tolerance against diols

All isolates from the 2,3-butanediol condition had mutations in the *relA* gene, the product of which is responsible for the synthesis of the stringent response alarmone ppGpp. This suggests that the toxicity of 2,3-butanediol induces an activation of stringent response leading to halted growth, and that an initial adaptive strategy is to prevent this response. One possible mechanism of stringent response activation would be potential membrane damage caused by 2,3-butadiol analoguously to membrane damage caused by ethanol stress[8]. Interestingly, the deletion of *relA* alone did not confer tolerance to 2,3-butanediol (Supplementary Figure 6), whereas the deletion of *sspA* (stringent activation protein A) resulted in high level of 2,3-butanediol tolerance.

We found that the growth inhibition by 2,3-butanediol could be alleviated by supplementation with methionine (data not shown). Additionally, at least one strain from each of the 2,3-butanediol populations had a mutation in the *metJ* gene, encoding a repressor of methionine synthesis. *metJ* deletions alone conferred relatively high level of tolerance to 2,3-butanediol (Supplementary Figure 6). Mutations in *metJ* are likely an adaptive response to increase intracellular methionine levels, to counter the inhibition by 2,3-butanediol. These results further support the notion that 2,3-butanediol stress has similarity to ethanol stress, where methionine supplementation also has been shown to reduce toxicity[9].

The only 2,3-butanediol evolved strain that had significantly increased endogenous production of 2,3-butanediol had a two unique mutations compared to two other strains from the same population (Supplementary Table 2): 1) a frameshift in the *acrB* gene encoding a subunit of the AcrAB-TolC multidrug efflux pump, and 2) a mobile element insertion in the *purT* gene. It is not clear why deletion of an efflux pump would be beneficial for production, but *acrB* deletions alone, and in combinations with a *metJ* deletion were shown to increase tolerance to 2,3-butanediol (Supplementary Figure 6). It has been previously found that acrA/acrB inactivations also improve isobutanol tolerance through an unknown mechanism[10].On the other hand *purT* deletion alone or in combination with *metJ* deletion was not found to be beneficial for 2,3-butanediol tolerance (Supplementary Figure 6).

As discussed in the main text, the majority of strains evolved on 1,2-propanediol were determined to be mutator strains. While the mutator phenotype meant that we could not include the strains in our overall analysis of the genetic landscape of tolerance, we can still determine if same gene targets that were mutated in 2,3-butanediol-evolved strains were also mutated in 1,2-propanediol-evolved strains. We found this to be the case for genes like *metJ, sspA* and *relA,* and thus we also tested the strains reconstructed based on 2,3-butanediol-evolved strains for tolerance of 1,2-propanediol. Again, *metJ* and *sspA* deletion strains grew significantly better than wild type strain, but in the 1,2-propanediol conditions, strains with combinations of more than one mutation did not grow better than these single deletions.

#### Evolved tolerance against diamines

Strains evolved to tolerate HMDA and putrescine had a high frequency of mutations in genes that have previously been implicated in osmotic stress resistance (*proV/X*, *nagA/C*)[11]. This is not surprising giving the high osmolarity of the final ALE condition. Deletion of *proV* alone conferred an increase in tolerance to both diamines with a much stronger effect on HMDA than putrescine tolerance. Other commonly mutated genes identified in diamine evolutions include *mreB*, encoding for an actin-like cell shape determining protein. A *mreB* deletion in combination with other deletions or genetic changes observed in strains that had *mreB* mutations conferred high levels of tolerance to both diamines (Supplementary Figure 6). In particular, strain having a *mreB* deletion together with a point mutation resulting in truncation of RpsG (S7 protein of S30 subunit of the ribosome) had high growth rates on both diamines. We tested many other combinations of mutations observed in specific diamine evolved strains (Supplementary Figure 6) and discovered several combinations that conferred high level of tolerance. However, it was difficult to determine a general mechanism of tolerance from this data.

#### Evolved tolerance against hexanoate

Mutations in *rpoA* (coding for RNA polymerase subunit alpha) were identified in almost all hexanoate-evolved isolates (most often clones had multiple unique mutations in *rpoA*), but these mutations could not be implemented in isolation in reconstructed strains. Out of the mutations tested, the combination of deletion with *ompR* (encoding for the response regulator of the EnvZ/OmpR two component system) together with either *bioAB* or *proP* gave most significant growth improvement on hexanoate. OmpR regulates the expression of membrane porins OmpF and OmpC as well as a number of other membrane-located genes[12]. Given that medium osmolarity is not particularly high in hexanoate evolution conditions, it is not likely that *ompR* mutations are in response to osmolarity changes, but rather relate to other membrane-related processes.

#### Evolved tolerance against octanoate

No consistent mutated genes were found across all octanoate-evolved strains, and the testing of some of the specific single deletion mutations also did not show significantly increased tolerance (Supplementary Figure 6). We also found no significant overlap between mutated genes in hexanoate and octanoate-evolved strains, so reconstructed hexanoate strains discussed above were not tested for tolerance of octanoate. Some of the octanoate-evolved strains had mutations in *mreB* gene, discussed above, as well as RNA polymerase subunit coding genes (*rpoA* and *rpoC)* indicating that morphological and broad regulatory changes play a role in tolerance. Laboratory evolution has previously been used to generate octanoate tolerating *E. coli* strains[13], but sequencing data from the previous work is not available.

#### Evolved tolerance against coumarate

Most coumaric-acid evolved strains had mutations in the termination factor *rho*. We were not able to implement the *rho* mutations detected individually in reconstructed strains. Additional interesting mutation targets discovered in the coumaric acid evolved strains were *mprA* and *nadR. mprA* encodes for a repressor of multidrug resistance pumps and deletion of it would result in upregulation of the pumps, which may allow pumping out coumarate. *nadR* encodes a multifunctional protein that acts both as a transcriptional repressor of transporters and NAD biosynthetic genes, and as a nicotinamide mononucleotide adenylyltransferase. *nadR* mutations are commonly seen in laboratory evolution studies (http://aledb.org and e.g. methanol utilization ALE[14]. Most of the *nadR* mutations we observe in coumarate-evolved strains are in the likely loss-of-function mutations and these mutations would result in derepression of NAD biosynthesis. The most tolerant reconstructed strain contained a triple deletion of *nadR, mprA* and *yhjK* (also known as *pdeK*) encoding for a putative c-di-GMP phosphodiesterase.

#### Evolved tolerance against n-butanol

We included n-butanol in the set of chemicals primarily to act as a control, since multiple previous studies have evolved *E. coli* to tolerate n-butanol[15–19]. Some of these studies have found similar genes commonly mutated to the ones we found (e.g. *marC* and *rob*) whereas many of the genes we found associated with n-butanol tolerance have not been previously observed (e.g. *yobF*). Our growth screening setup was not suitable for reliably determining growth rates in the presence of n-butanol due to significant evaporation, and hence we were not able to reliably verify whether individual mutations could confer tolerance to n-butanol (Supplementary Figure 6).

### References


1. Zhang, K., Woodruff, A. P., Xiong, M., Zhou, J. & Dhande, Y. K. A synthetic metabolic pathway for production of the platform chemical isobutyric acid. *ChemSusChem* **4**, 1068–1070 (2011).

2. Xu, Y. *et al.* Systematic metabolic engineering of Escherichia coli for high-yield production of fuel bio-chemical 2,3-butanediol. *Metab. Eng.* **23**, 22–33 (2014).

3. LaCroix, R. A., Palsson, B. O. & Feist, A. M. A model for designing adaptive laboratory evolution experiments. *Appl. Environ. Microbiol.* **83**, (2017).

4. Lennen, R. M. *et al.* Membrane stresses induced by overproduction of free fatty acids in Escherichia coli. *Appl. Environ. Microbiol.* **77**, 8114–8128 (2011).

5. Lennen, R. M. *et al.* Transient overexpression of DNA adenine methylase enables efficient and mobile genome engineering with reduced off-target effects. *Nucleic Acids Res.* **44**, 1–14 (2015).

6. Deatherage, D. E. & Barrick, J. E. Identification of mutations in laboratory evolved microbes from next-generation sequencing data using breseq. *Methods Mol. Biol.* **1151**, 165–188 (2014).

7. Phaneuf, P. V, Gosting, D., Palsson, B. O. & Feist, A. M. ALEdb 1.0: A Database of Mutations from Adaptive Laboratory Evolution Experimentation. *Nucleic Acids Res.* 1–8 (2018). doi:10.1093/nar/gky983

8. Cao, H. *et al.* Systems-level understanding of ethanol-induced stresses and adaptation in E. coli. *Sci. Rep.* **7**, 44150 (2017).

9. Haft, R. J. F. *et al.* Correcting direct effects of ethanol on translation and transcription machinery confers ethanol tolerance in bacteria. *Proc. Natl. Acad. Sci.* **111**, E2576--E2585 (2014).

10. Lee, S. Y. & Kim, H. U. Systems strategies for developing industrial microbial strains. *Nat.*

*Biotechnol.* **33**, 1061–1072 (2015).

11. Winkler, J. D., Garcia, C., Olson, M., Callaway, E. & Kao, K. C. Evolved osmotolerant escherichia coli mutants frequently exhibit defective N-acetylglucosamine catabolism and point mutations in cell shape-regulating protein MreB. *Appl. Environ. Microbiol.* **80**, 3729–3740 (2014).

12. Seo, S. W. *et al.* Revealing genome-scale transcriptional regulatory landscape of OmpR highlights its expanded regulatory roles under osmotic stress in Escherichia coli K-12 MG1655. *Sci. Rep.* **7**, 1–10 (2017).

13. Royce, L. A. *et al.* Evolution for exogenous octanoic acid tolerance improves carboxylic acid production and membrane integrity. *Metab. Eng.* **29**, 180–188 (2015).

14. Meyer, F. *et al.* Methanol-essential growth of Escherichia coli. *Nat. Commun.* **9**, (2018).

15. Reyes, L. H., Abdelaal, A. S. & Kao, K. C. Genetic determinants for n-butanol tolerance in evolved escherichia coli mutants: Cross adaptation and antagonistic pleiotropy between n-butanol and other stressors. *Appl. Environ. Microbiol.* **79**, 5313–5320 (2013).

16. Reyes, L. H., Almario, M. P., Winkler, J., Orozco, M. M. & Kao, K. C. Visualizing evolution in real time to determine the molecular mechanisms of n-butanol tolerance in Escherichia coli. *Metab. Eng.* **14**, 579–590 (2012).

17. Zhu, L., Li, Y. & Cai, Z. Development of a stress-induced mutagenesis module for autonomous adaptive evolution of Escherichia coli to improve its stress tolerance. *Biotechnol. Biofuels* **8**, 1–10 (2015).

18. Zorraquino, V., Kim, M., Rai, N. & Tagkopoulos, I. The genetic and transcriptional basis of short and long term adaptation across multiple stresses in Escherichia coli. *Mol. Biol. Evol.* **34**, 707–717 (2017).

19. Horinouchi, T. *et al.* Prediction of Cross-resistance and Collateral Sensitivity by Gene Expression profiles and Genomic Mutations. *Sci. Rep.* **7**, 1–11 (2017).

### Supplementary Tables

*Supplementary Table 1: Concentrations of each chemical used during ALE and for growth screening. All concentrations are g/L.*

| | INITIAL ALE CONCENTRATION | FINAL ALE CONCENTRATION | ISOLATE SCREENING CONCENTRATION | CROSS-TOLERANCE SCREENING CONCENTRATION |
|---|---|---|---|---|
| **1,2-PROPANEDIOL** | 52 | 83 | 83 | 62 |
| **2,3-BUTANEDIOL** | 49 | 79 | 69 | 59 |
| **HEXAMETHYLENEDIAMINE** | 20 | 38 | 38 | 32 |
| **PUTRESCINE** | 20 | 38 | 38 | 32 |
| **GLUTARATE** | 20 | 47.5 | 47.5 | 40 |
| **ADIPATE** | 25 | 50 | 50 | 45 |
| **HEXANOATE** | 2 | 7.5 | 5 | 3 |
| **OCTANOATE** | 3.5 | 10 | 8 | 8 |
| **ISOBUTYRATE** | 3 | 12.5 | 12.5 | 7.5 |
| **COUMARATE** | 4 | 20 | 10 | 7.5 |
| **BUTANOL** | 5.7 | 16.2 | 11.34 | 11.34 |

*Supplementary Table 2: Summary of the mutations and mutated genes in all the non-mutator strains. The mutation names denote the type, location and change of the mutations.*

| | Genes | Mutations |
|---|---|---|
| 12PD4-6 | relA, metJ, yeaR, sspA, rpsA | SNP-3377240-G, SNP-4128078-G, SNP-2912634-G, SNP-962923-T, MOB-1879829-Δ1-: |
| 12PD6-3 | lrhA, rpoA, fabR, dusA, yfgF | SNP-4161155-A, SNP-4261586-C, SNP-3440194-A, MOB-2628616-IS2-5, MOB-2406831-IS2-5 |
| 12PD6-9 | rpoA, fabR, yfgF, ypjA | SNP-3440194-A, MOB-2628616-IS2-5, SNP-2780609-C, SNP-4161155-A |
| 23BD1-6 | gabP, nanK, rnb, metJ, elfA, rpoB, purT, relA | MOB-998193-IS5-4, SNP-4182583-T, SNP-2794550-G, SNP-1931977-G, SNP-3369969-A, DEL-2911491-7528, SNP-4128380-A, MOB-1347480-IS5-4 |
| 23BD1-9 | gabP, nanK, rnb, metJ, elfA, rpoB, purT, relA | MOB-998193-IS5-4, SNP-4182583-T, SNP-2794550-G, SNP-1931977-G, SNP-3369969-A, DEL-2911491-7528, SNP-4128380-A, MOB-1347480-IS5-4 |
| 23BD2-4 | nanK, uspC, metJ, rpoC, rpsA, gtrS, relA | SNP-1979639-C, MOB-4128293-IS5-4, SNP-4186152-G, SNP-3369969-A, SNP-962056-T, MOB-580116-IS5-4, SNP-2470411-G, DEL-2911491-7528 |
| 23BD2-7 | metJ, rpoC, nanK, relA, rpsA | MOB-4128293-IS5-4, MOB-1096841-IS2-5, SNP-4186152-G, SNP-3369969-A, SNP-962056-T, MOB-580116-IS5-4, DEL-2911491-7528 |
| 23BD2-9 | metJ, rpoC, nanK, relA, rpsA | MOB-4128293-IS5-4, SNP-4186152-G, SNP-3369969-A, SNP-962056-T, MOB-580116-IS5-4, DEL-2911491-7528 |
| 23BD4-3 | yeaR, mprA, rnb, fadB, umuD, metJ, rpoC, purT, relA | SNP-1347775-A, SNP-4128361-C, SNP-2810756-T, SNP-2913536-T, DEL-4187816-15, SNP-1931977-G, MOB-1879829-Δ1-:, SNP-4031019-A, SNP-1230727-A |
| 23BD4-4 | yeaR, umuD, rpoC, metJ, purT, relA | SNP-4128361-C, SNP-1931977-G, MOB-1879829-Δ1-:, SNP-2913536-T, DEL-4187816-15, SNP-1230727-A |
| 23BD4-7 | rpoC, nanK, metJ, relA | SNP-4186274-T, SNP-3369969-A, SNP-4128169-G, DEL-2911491-7528 |
| 23BD5-1 | yeaR, spoT, umuD, metJ, rpoC, purT, relA | SNP-4128361-C, MOB-1879829-Δ1-:, SNP-3823036-C, SNP-2913536-T, DEL-4187816-15, SNP-1931977-G, SNP-1230727-A |
| 23BD5-7 | ybeT, nanK, ybhP, ydhK, rpoC, metJ, zntR, relA | SNP-4186274-T, SNP-1722386-T, SNP-4128379-T, SNP-824028-G, SNP-3369969-A, SNP-3438773-G, DEL-2911491-7528, SNP-679090-G |
| 23BD5-10 | rpoC, nanK, metJ, relA | SNP-4186274-T, SNP-3369969-A, SNP-4128169-G, DEL-2911491-7528 |
| 23BD6-1 | essD, rpoC, metJ, nusG, purT, relA | DEL-575786-3027, DEL-2912618-10, SNP-4185573-G, SNP-1930993-G, SNP-4128250-C, SNP-4178172-G |
| 23BD7-4 | treR, nanK, elfD, tolC, metJ, rpoC, yhjA, flu, purT, relA | SNP-4128386-C, MOB-3668878-IS2-5, SNP-4186152-G, MOB-4466841-IS5-4, SNP-1931668-G, SNP-3369969-A, DEL-2911491-7528, SNP-3178128-G, SNP-2073463-A, MOB-998719-IS2-5 |

| | | |
|---|---|---|
| 23BD7-5 | acrB, nanK, elfD, metJ, rpoC, flu, purT, relA | SNP-4128386-C, SNP-4186152-G, SNP-3369969-A, MOB-998719-IS2-5, DEL-2911491-7528, INS-484102-AT, SNP-2073463-A, MOB-1931499-IS5-4 |
| 23BD7-7 | treR, nanK, elfD, tolC, metJ, rpoC, yhjA, flu, purT, relA | SNP-4128386-C, MOB-3668878-IS2-5, SNP-4186152-G, MOB-4466841-IS5-4, SNP-1931668-G, SNP-3369969-A, DEL-2911491-7528, SNP-3178128-G, SNP-2073463-A, MOB-998719-IS2-5 |
| 23BD8-2 | ygaH, iscR, relA, rpoB, pyrE, lon | SNP-2661816-G, SNP-2913641-T, SNP-4184579-G, DEL-3815810-1, SNP-2810459-C, SNP-461034-G |
| 23BD8-7 | ygaH, iscR, metJ, lacZ, rpoB, pyrE, relA | DEL-3815810-1, DEL-365742-1, SNP-2661816-G, SNP-4128212-G, SNP-2810459-C, SNP-2913641-T, SNP-4184579-G |
| ADIP1-1 | sspA, kgtP, gltP, ybjL, proV, yicC, uvrB, pyrE | SNP-814029-G, DEL-3377068-21, MOB-2725207-IS1-9, SNP-3815823-A, DEL-2804648-38, SNP-4294366-T, DEL-889569-1, SNP-3816848-T |
| ADIP1-9 | allD, sspA, kgtP, ligA, gltP, ybjL, proV, yicC, pyrE | DEL-3377068-21, SNP-2530235-T, MOB-2725207-IS1-9, SNP-3815823-A, SNP-546309-T, DEL-2804648-38, SNP-4294366-T, DEL-889569-1, SNP-3816848-T |
| ADIP2-5 | lacY, rph, nagC, kgtP | SNP-362830-A, MOB-700529-IS1-9, DEL-3815884-2, SNP-2725613-A |
| ADIP2-6 | lacY, rph, nagC, kgtP | MOB-700529-IS1-9, SNP-362830-A, SNP-2725613-A, DEL-3815884-2 |
| ADIP2-10 | ydcD, pyrE, nagC, kgtP | SNP-1530007-A, MOB-700628-IS5-4, DEL-3815810-1, SNP-2725613-A |
| ADIP3-2 | yfgO, sspA, kgtP, ybjL, proV, yicC, pyrE | DEL-3377068-21, MOB-2725207-IS1-9, SNP-3815823-A, DEL-2804648-38, SNP-2614996-G, SNP-3816848-T, MOB-889534-IS5-4 |
| ADIP3-4 | yeaR, kgtP, sspA, ybjL, yhiL, proV, yicC, mltD, pyrE | DEL-3377068-21, INS-233963-GT, MOB-2725207-IS1-9, SNP-3815823-A, MOB-3633911-IS5-4, DEL-2804648-38, MOB-1879829-Δ1-:, SNP-3816848-T, MOB-889534-IS5-4 |
| ADIP3-8 | sspA, kgtP, ybjL, proV, yicC, pyrE | DEL-3377068-21, MOB-2725207-IS1-9, SNP-3815823-A, DEL-2804648-38, SNP-3816848-T, MOB-889534-IS5-4 |
| ADIP4-8 | purL, kgtP, malQ, ybjL, yphC, pdxJ, srmB, nagA, rnt, metL | DEL-4130167-462, SNP-1728708-G, SNP-2713302-T, MOB-889488-IS1-9, SNP-2675452-C, SNP-2724590-T, SNP-3548179-T, SNP-2701175-C, SNP-2693818-G, INS-702444-GCATAACGCGCACGCCCTGTTTCATCAGCTCATCGC |
| ADIP6-3 | spoT, kgtP, ubiE, proQ, nagC, ybjL, proV | SNP-3823700-T, SNP-700928-A, DEL-2804864-13, SNP-1915297-A, SNP-2725155-T, MOB-889534-IS5-4, SNP-4019173-G |
| ADIP6-9 | spoT, kgtP, proQ, nagC, ybjL, proV, icd, ycjG | SNP-3823700-T, SNP-1196319-A, SNP-700928-A, SNP-1915297-A, SNP-2725155-T, MOB-889534-IS5-4, SNP-1389396-C, DEL-2804835-7 |
| ADIP6-10 | spoT, kgtP, proQ, nagC, ybjL, proV | SNP-2805493-T, SNP-3823700-T, SNP-700928-A, SNP-1915297-A, SNP-2725155-T, MOB-889534-IS5-4 |
| ADIP7-2 | kgtP, yagL, rpoS, yneO, hns, pstS | SNP-2725329-A, SNP-293574-G, MOB-1293038-IS1-9, MOB-1598223-IS5-4, DEL-2867359-9, INS-3911001-TTTC |
| ADIP7-5 | ybjL, idnR, proV, sspA, kgtP | MOB-889540-IS5-4, SNP-3377240-G, DEL-2725643-1, MOB-4490689-IS1-9, DEL-2804864-13 |
| ADIP8-3 | pstS, yehD, hns, kgtP | MOB-1293015-IS1-8, SNP-2724588-T, DEL-2192452-1, MOB-3911563-IS1-9 |
| ADIP8-7 | pstS, yehD, hns, kgtP | MOB-1293015-IS1-8, SNP-2724588-T, MOB-3911563-IS1-9, DEL-2192452-1 |
| ADIP8-10 | pstS, insI1, yehD, hns, kgtP | MOB-1293015-IS1-8, MOB-280003-IS5-4, SNP-2724588-T, DEL-2192452-1, MOB-3911563-IS1-9 |
| BUT1-2 | tqsA, adeP, manY, rob, cspC, pyrE | DEL-3815810-1, SNP-3895843-C, DEL-1907330-2, SNP-1903497-C, SNP-4635114-C, MOB-1673883-IS5-4 |
| BUT1-3 | manY, pyrE, tqsA, cspC, rob | SNP-4635114-C, SNP-1903497-C, MOB-1673883-IS5-4, DEL-1907330-2, DEL-3815810-1 |
| BUT1-5 | tqsA, manY, yobF, ycaN, rob, pyrE | MOB-1907448-IS5-4, DEL-3815810-1, SNP-1674448-A, INS-4635196-G, SNP-1903497-C, SNP-949006-G |
| BUT2-9 | hfq, marC, rob | SNP-1618698-A, SNP-4400407-G, DEL-3815859-82, INS-4634895-A |
| BUT3-3 | manY, yobF, leuA, marC, pyrE | DEL-1618281-1, SNP-1903497-C, MOB-1907448-IS5-4, DEL-3815810-1, SNP-82594-G |
| BUT3-6 | tqsA, manY, yobF, pheU, marC, pyrE | INS-4362600-C, MOB-1907448-IS5-4, DEL-3815810-1, SNP-1903497-C, MOB-1673883-IS5-4, DEL-1618281-1 |
| BUT3-7 | tqsA, manY, yobF, pheU, marC, pyrE | INS-4362600-C, MOB-1907448-IS5-4, DEL-3815810-1, SNP-1903497-C, MOB-1673883-IS5-4, DEL-1618281-1 |
| BUT4-4 | manY, marC, rob, glmU, cspC, pyrE | MOB-1907343-IS1-8, MOB-1618666-IS2-5, DEL-3815810-1, SNP-4635159-T, SNP-1903497-C, SNP-3915089-T |
| BUT4-7 | manY, marC, rob, glmU, cspC, pyrE, sapA | MOB-1907343-IS1-8, MOB-1618666-IS2-5, DEL-3815810-1, SNP-4635159-T, SNP-1356672-G, SNP-1903497-C, SNP-3915089-T |
| BUT4-9 | manY, marC, rob, glmU, cspC, pyrE | MOB-1907343-IS1-8, MOB-1618666-IS2-5, DEL-3815810-1, SNP-4635159-T, SNP-1903497-C, SNP-3915089-T |
| BUT5-2 | manY, marC, mppA, rob, glmU, cspC, pyrE | MOB-1907343-IS1-8, DEL-3815810-1, SNP-4635159-T, DEL-1394081-2, SNP-1903497-C, MOB-1618850-IS5-4, SNP-3915089-T |
| BUT5-3 | manY, marC, rob, glmU, cspC, pyrE | MOB-1907343-IS1-8, MOB-1618666-IS2-5, DEL-3815810-1, SNP-4635159-T, SNP-1903497-C, SNP-3915089-T |
| BUT6-1 | manY, yobF, marC, pyrE, rob | SNP-4635243-T, SNP-1903497-C, MOB-1907448-IS5-4, DEL-3815810-1, DEL-1606886-11558 |
| BUT6-3 | manY, yobF, marC, pyrE, rob | DEL-1606886-11558, SNP-1903497-C, MOB-1907448-IS5-4, SNP-4635243-T, DEL-3815810-1 |

| | | |
|---|---|---|
| BUT6-8 | manY, yobF, marC, pyrE, rob | SNP-4635243-T, SNP-1903497-C, MOB-1907448-IS5-4, DEL-3815810-1, DEL-1606886-11558 |
| BUT7-6 | manY, yobF, pyrE, rob, mppA | MOB-1907611-IS5-4, SNP-4635203-T, SNP-1903497-C, DEL-3815810-1, DEL-1392752-3292 |
| BUT7-7 | manY, yobF, pyrE, rob, mppA | MOB-1907611-IS5-4, SNP-4635203-T, SNP-1903497-C, DEL-3815810-1, DEL-1392752-3292 |
| BUT7-9 | manY, yobF, pyrE, rob, mppA | MOB-1907611-IS5-4, SNP-4635203-T, SNP-1903497-C, DEL-3815810-1, DEL-1392752-3292 |
| BUT9-7 | pyrE, manY, rob, marC, rraA | DEL-4119238-18, INS-1618379-C, SNP-4635048-C, DEL-3815810-1, SNP-1903497-C |
| BUT9-10 | rraA, manY, marC, rob, pyrE, otsB | INS-1618379-C, SNP-4635048-C, DEL-3815810-1, SNP-1903497-C, SNP-1981720-A, DEL-4119238-18 |
| COUM1-2 | rho, fimD, ycfQ, sapC, rpoC, polB, nadR | SNP-3966727-T, SNP-4627958-T, DEL-4546637-1, SNP-1168483-T, SNP-4185540-T, SNP-1354284-A, SNP-64352-C |
| COUM2-3 | rho, atpI, murC, sapF, rpoC, nadR | SNP-1352163-A, DEL-102228-1, MOB-3922629-IS5-4, DEL-4627451-124, SNP-3966751-T, SNP-4185540-T |
| COUM2-4 | rho, atpI, murC, ccmA, sapF, rhaT, rpoC, nadR, yecT | SNP-3966751-T, DEL-102228-1, MOB-2297586-IS5-4, DEL-4627451-124, SNP-4185540-T, SNP-4099695-A, SNP-1352163-A, MOB-1961829-IS5-4, MOB-3922629-IS5-4 |
| COUM2-7 | rho, atpI, murC, sapF, rpoC, nadR | SNP-3966751-T, DEL-102228-1, DEL-4627451-124, SNP-4185540-T, SNP-1352163-A, MOB-3922629-IS5-4 |
| COUM3-1 | rho, atpI, mprA, dacA, manY, rpoB, hns, pyrE | SNP-663746-T, DEL-3815810-1, DEL-2810080-1165, MOB-1293196-IS5-4, SNP-3966727-T, DEL-260217-13738, SNP-3922483-A, SNP-4183802-G, SNP-1903497-C |
| COUM3-9 | rho, atpI, mprA, dacA, manY, rpoB, hns, pyrE | SNP-663746-T, DEL-3815810-1, DEL-2810080-1165, MOB-1293196-IS5-4, SNP-3966727-T, SNP-4183802-G, SNP-3922483-A, SNP-1903497-C |
| COUM3-10 | rho, atpI, mprA, dacA, manY, rpoB, hns, pyrE | SNP-663746-T, DEL-3815810-1, DEL-2810080-1165, MOB-1293196-IS5-4, SNP-3966727-T, SNP-4183802-G, SNP-3922483-A, SNP-1903497-C |
| COUM4-2 | epmB, rpoA, rpsG, nadR, dcd | SNP-4375431-C, SNP-3473615-T, SNP-2141832-T, SNP-3440924-G, SNP-4627567-T |
| COUM4-5 | rpsG, dcd, yfiN, ompN, nadR, rpoA | SNP-3473615-T, SNP-2141832-T, SNP-3440924-G, SNP-2743181-G, SNP-4627567-T, SNP-1436746-C |
| COUM4-10 | rpoA, rpsG, nadR, dcd | SNP-3473615-T, SNP-2141832-T, SNP-3440924-G, SNP-4627567-T |
| COUM5-3 | rho, atpI, sapF, yphF, ypjC, glnG, rpoC, mrdA, nadR | SNP-3966751-T, MOB-2784452-IS5-4, MOB-2678755-IS5-4, DEL-4054531-104, SNP-667158-T, SNP-4185540-T, SNP-1352163-A, DEL-4628232-1, MOB-3922629-IS5-4 |
| COUM5-5 | rho, sapF, ypjC, rpoC, mrdA, nadR | SNP-3966751-T, MOB-2784452-IS5-4, SNP-667158-T, SNP-4185540-T, DEL-4628214-1, SNP-1352163-A |
| COUM5-8 | focA, rho, atpI, sapF, ypjC, tufA, rpoC, mrdA, ydiJ | SNP-3966751-T, MOB-2784452-IS5-4, SNP-954638-T, SNP-1768309-T, SNP-667158-T, SNP-4185540-T, DEL-3471319-1, SNP-1352163-A, INS-3922570-TAG |
| COUM6-2 | rho, yjiP, manY, rnb, yhgE, nusA, pyrE | MOB-1347892-IS5-4, DEL-3815810-1, SNP-3966727-T, SNP-4569172-A, SNP-3317438-A, SNP-1903497-C, SNP-3530902-G |
| COUM6-5 | rho, yjiP, manY, rnb, yhgE, nusA, pyrE | MOB-1347892-IS5-4, DEL-3815810-1, SNP-3966727-T, SNP-4569172-A, SNP-3317438-A, SNP-1903497-C, SNP-3530902-G |
| COUM6-9 | rho, yjiP, manY, rnb, yhgE, nusA, fimC, pyrE | DEL-3815810-1, SNP-3530902-G, MOB-4544671-+G-S5, MOB-1347892-IS5-4, SNP-3966727-T, SNP-4569172-A, SNP-3317438-A, SNP-1903497-C |
| COUM7-5 | rho, mprA, ypjA, manY, prlF, rpoB, ydjH, pyrE | SNP-4183814-A, DEL-3815810-1, SNP-3966727-T, DEL-2810804-1, MOB-1856052-IS5-4, SNP-1903497-C, INS-3277273-TTCAACA, MOB-2782626-IS5-4 |
| COUM7-6 | rho, mprA, ypjA, manY, mgrB, rpoB, pyrE | MOB-1908812-IS5-4, SNP-4183814-A, DEL-3815810-1, SNP-3966727-T, DEL-2810804-1, SNP-1903497-C, MOB-2782626-IS5-4 |
| COUM8-1 | manY, thrA, pyrE, mprA, yhjK | DEL-3815810-1, DEL-3683736-1, SNP-1903497-C, SNP-2374-T, DEL-2801966-11843 |
| COUM8-6 | manY, rho, pyrE, mprA, yhjK | INS-3966718-GAT, SNP-1903497-C, DEL-3685181-273, DEL-3815810-1, DEL-2801966-11843 |
| GLUT1-3 | spoT, kgtP, rnb, nagC, rpoC, ydfI | SNP-4186605-C, SNP-1630841-A, SNP-3823664-C, MOB-700614-IS1-9, DEL-2725672-1, SNP-1347104-T |
| GLUT1-9 | rpoC, rnb, spoT, nagC, kgtP | SNP-3823664-C, MOB-700614-IS1-9, DEL-2725672-1, SNP-4186605-C, SNP-1347104-T |
| GLUT1-10 | yiaT, hofM, spoT, insG, sspA, kgtP, proV, greA | SNP-3522182-A, INS-3377241-AGCTCACGATCCACCAGGGTC, INS-2805532-T, MOB-3328463-IS4-11, SNP-3823664-C, MOB-3751884-IS5-4, DEL-2725672-1 |
| GLUT2-1 | spoT, kgtP, tomB, ygjP, proV, nagA | SNP-3823751-A, MOB-701614-IS1-9, DEL-2725643-1, SNP-3236414-C, DEL-2804864-13, SNP-481075-G |
| GLUT2-9 | nagA, spoT, rnb, kgtP | SNP-3823751-A, MOB-701614-IS1-9, DEL-2725643-1, DEL-1347882-1 |
| GLUT2-10 | rspA, spoT, kgtP, rpoC, proV, nagA | SNP-1654069-C, SNP-4186605-C, SNP-3823751-A, MOB-701614-IS1-9, DEL-2725643-1, DEL-2804864-13 |
| GLUT3-5 | rpoC, spoT, kgtP | SNP-3823770-T, SNP-2724971-C, SNP-4186605-C |
| GLUT3-7 | rpoC, spoT, kgtP | SNP-3823770-T, SNP-2724971-C, SNP-4186605-C |
| GLUT3-9 | spoT, kgtP, wzzE, ssuA, rclB, rnt | SNP-318484-T, SNP-996768-A, SNP-1728882-C, INS-2725518-C, INS-3969051-G, SNP-3823759-C |
| GLUT4-1 | nagC, proX, spoT, kgtP | SNP-3824137-T, SNP-2724611-A, DEL-2807199-8, MOB-701188-IS1-9 |

| | | |
|---|---|---|
| GLUT4-4 | proX, spoT, nagC, kgtP | SNP-2390019-A, MOB-3195220-IS186-4, DEL-2807199-8, MOB-701188-IS1-9, SNP-3824137-T, SNP-2724611-A |
| GLUT4-10 | csiD, rpoC, spoT, kgtP | MOB-2788702-IS5-4, SNP-2724971-C, SNP-4186605-C, SNP-3823751-T |
| GLUT5-4 | ytfR, spoT, sspA, kgtP, yagU, rpoB | DEL-3377359-18, DEL-2725375-1, SNP-3823106-T, DEL-303121-1, SNP-4451123-A, SNP-4181852-C |
| GLUT5-5 | ytfR, spoT, rpoB, sspA, kgtP | MOB-3377491-IS2-5, SNP-4451123-A, DEL-2725375-1, SNP-3823106-T, SNP-4181852-C |
| GLUT5-9 | ytfR, spoT, rpoB, sspA, kgtP | MOB-3377491-IS2-5, SNP-4451123-A, DEL-2725375-1, SNP-3823106-T, SNP-4181852-C |
| GLUT6-4 | kgtP, spoT, yfjL, nagC, cspE | SNP-3823105-A, DEL-657215-25, MOB-700680-IS1-9, DEL-2765456-8, SNP-2725370-C |
| GLUT6-5 | nagC, spoT, yfjL, kgtP, cspE | SNP-2725370-C, SNP-3823105-A, DEL-657215-25, DEL-2765456-8, MOB-700680-IS1-9 |
| GLUT6-10 | hcaD, nagA, spoT, rnt, kgtP | INS-2724732-AAAAGC, MOB-701889-IS1-9, SNP-3823139-A, SNP-1728425-A, DEL-2672981-6 |
| GLUT7-2 | spoT, rnt, nagC, kgtP | SNP-701396-A, SNP-1728926-C, SNP-2724848-A, DEL-3824201-6 |
| GLUT7-6 | nohQ, spoT, rnt, kgtP | SNP-2724848-A, SNP-1636300-G, SNP-1728926-C, DEL-3824201-6 |
| GLUT7-7 | kgtP, rlmI, yhfA, ravA, spoT, dkgA, rplM, nagC, rrlA, rrlC, uvrD, ybeF, rsmC, fruB, yhiL, tdcD, yfdC, lldR, yliE, rnt, aceK, roxA, yhfX | SNP-443040-T, DEL-3943892-1, DEL-1029739-1, SNP-3823724-T, SNP-2262665-A, DEL-2465722-1, SNP-3263200-A, DEL-701381-1, SNP-660573-C, SNP-4219696-T, SNP-3779384-G, SNP-4037513-C, INS-3931183-G, DEL-2725672-1, SNP-3510180-C, SNP-874067-A, SNP-1728884-A, SNP-3378539-G, SNP-3485967-G, SNP-3634152-C, SNP-3156603-C, SNP-4607712-T, SNP-1187352-A, DEL-3999387-1 |
| GLUT8-5 | rpoC, polB, proV, mprA, kgtP | SNP-2725818-G, DEL-2804864-13, MOB-2810987-IS1-9, SNP-4185540-T, SNP-64352-C |
| GLUT8-6 | kgtP, sapC, rpoC, polB, proV, nagA | DEL-2804864-13, SNP-2725232-A, SNP-1354284-A, SNP-4185540-T, INS-702338-T, SNP-64352-C |
| GLUT8-9 | kgtP, yobF, sapC, sdaC, rpoC, proV, polB, lit | MOB-1907448-IS5-4, DEL-2804926-1, INS-1198505-AATGATGA, DEL-2725209-9, SNP-4185540-T, SNP-2927703-A, SNP-1354284-A, SNP-64352-C |
| HEXA1-1 | ptrA, rpoA, rpoC, bioB, sapB | SNP-809340-G, SNP-3440378-T, SNP-3440212-A, SNP-4185540-T, MOB-2957831-IS5-4, SNP-1354687-A |
| HEXA1-4 | opgH, rpoA, rpoC, bioB, sapB | SNP-809340-G, SNP-3440378-T, DEL-1112435-5, SNP-3440212-A, SNP-4185540-T, SNP-1354687-A |
| HEXA1-5 | sapB, rpoA, rpoC, bioB | SNP-809340-G, SNP-3440378-T, SNP-1354687-A, SNP-4185540-T, SNP-3440212-A |
| HEXA2-3 | pykF, ompR, mdtK, prpE | SNP-353944-A, SNP-1743611-A, DEL-3536285-1, SNP-1756622-A |
| HEXA2-9 | ompR, mdtK, prpE | SNP-1743611-A, SNP-353944-A, DEL-3536285-1 |
| HEXA2-10 | rpoA, mdtK | SNP-3440212-A, SNP-3440929-A, DEL-1744016-757 |
| HEXA3-1 | rpoC, rpoA, yfjL, mdtK, sapA | SNP-1356317-T, SNP-3440378-T, MOB-1743880-IS1-9, SNP-3440212-A, SNP-2764096-T, SNP-4185540-T |
| HEXA3-7 | mdtK, rpoA, rpoC, yfjL | SNP-3440212-A, SNP-3440378-T, SNP-2764096-T, MOB-1743880-IS1-9, SNP-4185540-T |
| HEXA3-9 | mdtK, rpoA, rpoC, yfjL | SNP-3440212-A, SNP-3440378-T, SNP-2764096-T, MOB-1743880-IS1-9, SNP-4185540-T |
| HEXA4-4 | rpoA, ompR, proQ, glxK, hns | MOB-1293196-IS5-4, MOB-542938-IS5-4, DEL-1915293-5, INS-3536332-T, SNP-3440923-T |
| HEXA4-7 | dosP, rpoA, ompR, proQ, hns | MOB-1293196-IS5-4, INS-3536332-T, DEL-1915293-5, SNP-1564102-G, SNP-3440923-T |
| HEXA4-10 | rpoA, ompR, proQ, hns | MOB-1293196-IS5-4, INS-3536332-T, SNP-1915353-C, SNP-3440923-T |
| HEXA6-5 | sapB, rpoA, mdtK | SNP-3440212-C, DEL-1744675-1, SNP-1354687-A, INS-3440937-CGCTCT |
| HEXA6-6 | rpoA, mdtK | SNP-3440212-C, DEL-1744675-1, INS-3440937-CGCTCT |
| HEXA6-7 | rhmD, rpoA, mdtK, sapB | SNP-3440212-C, SNP-2360829-A, DEL-1744675-1, SNP-1354687-A, INS-3440937-CGCTCT |
| HEXA6-9 | mdtK, rpoA, rsd, sapB | SNP-3440212-C, DEL-1744675-1, INS-3440937-CGCTCT, SNP-1354687-A |
| HEXA7-2 | ompC, emrY, rsfS, sfmF, rpoB, hns, rpoA | MOB-563324-IS1-9, SNP-2481325-A, SNP-4182358-T, MOB-2312877-IS5-4, MOB-1293124-IS5-4, SNP-3440923-T, DEL-668970-1 |
| HEXA8-1 | ompC, yedP, sapB, cydA, barA, murG, rpoA | SNP-99891-T, SNP-1354761-T, INS-2025248-TC, MOB-771258-IS5-4, MOB-2312877-IS5-4, DEL-2916392-12, SNP-3440923-T |
| HEXA8-2 | ompC, yedP, sapB, cydA, murG, rpoA | SNP-99891-T, MOB-771306-IS5-4, SNP-1354761-T, INS-2025248-TC, MOB-2312877-IS5-4, SNP-3440923-T |
| HEXA8-5 | ompC, yedP, sapB, murG, rpoC, rpoA | SNP-99891-T, SNP-1354761-T, INS-2025248-TC, MOB-2312877-IS5-4, SNP-3440923-T, SNP-4187619-T |
| HMDA1-10 | purL, rph, spoT, lexA, proV, rpoB, rpsA | SNP-2694102-A, SNP-962939-A, SNP-3816611-A, DEL-2804864-13, SNP-4257602-T, SNP-4181786-T, SNP-3823025-A |
| HMDA2-1 | ptsP, proV, pyrE, rpsA | MOB-2804836-IS1-9, DEL-2968163-1, DEL-3815808-1, SUB-963273- |
| HMDA2-8 | ptsP, proV, pyrE, rpsA | MOB-2804836-IS1-9, SUB-963273-, DEL-2968163-1, DEL-3815808-1 |

| | | |
|---|---|---|
| HMDA3-4 | rpoC, nagA, pyrE, kup | SNP-3933122-A, INS-702597-G, DEL-3815810-1, SNP-4188767-T |
| HMDA3-5 | nagC, ygeG, kup, ygbT, pyrE, ybeX | SNP-3933122-A, SNP-2879763-A, DEL-3815810-1, DEL-691774-12, SNP-701405-A, SNP-2991218-G |
| HMDA3-6 | gatY, rpoC, nagA, pyrE, kup | SNP-3933122-A, DEL-3815810-1, MOB-2177307-IS1-9, INS-702597-G, SNP-4188767-T |
| HMDA5-4 | ptsP, ampC, pnp, pyrE, nagC | SNP-2966573-G, DEL-3815810-1, SNP-3310266-A, SNP-4378331-G, MOB-700602-IS1-9 |
| HMDA5-5 | ptsP, pyrE, nagC, ybeX | SNP-2966573-G, DEL-3815810-1, MOB-700602-IS1-9, SNP-691321-T |
| HMDA5-10 | ptsP, pstB, pyrE, pepA, stpA | SNP-2966573-G, DEL-3815810-1, MOB-2798597-IS1-9, SNP-4485639-C, SNP-3908248-T |
| HMDA7-1 | rpsG, wbbK, sspA, nusA | SNP-3377173-C, DEL-2104077-1, SNP-3473612-C, SNP-3317072-C |
| HMDA7-7 | rpsG, sspA, nusA | SNP-3377173-C, SNP-3473612-C, SNP-3317072-C |
| HMDA7-10 | rpsG, wbbK, sspA, nusA | SNP-3377173-C, DEL-2104077-1, SNP-3473612-C, SNP-3317072-C |
| HMDA8-5 | mdtK, xapR, nagC, proV, cynR, pyrE, lhr, rnt | SNP-1728512-C, DEL-3815808-1, SNP-1732811-T, DEL-2804864-13, SNP-2522653-A, SNP-358399-G, SNP-700980-C, SNP-1744313-A |
| HMDA8-9 | mdtK, nagC, proV, pyrE, lhr, rnt | SNP-1728512-C, DEL-3815808-1, SNP-1732811-T, DEL-2804864-13, SNP-700980-C, SNP-1744313-A |
| HMDA8-10 | mpl, mdtK, nagC, proV, pyrE, lhr, rnt | SNP-1728512-C, DEL-3815808-1, SNP-1732811-T, DEL-2804864-13, DEL-4457113-4, SNP-700980-C, SNP-1744313-A |
| IBUA1-7 | ptsP, pykF, insA, rpoB | SNP-20771-A, MOB-1755755-IS5-4, MOB-2967576-IS5-4, SNP-4183097-T |
| IBUA1-9 | yedV, pykF, rlmE, rpoB, cheR | DEL-255591-18364, SNP-4182938-C, SNP-1969313-A, SNP-2037332-T, MOB-3327665-IS5-4, SNP-1756637-C |
| IBUA2-1 | rpsC, yobF, yijD, bglF, sapD, rpoC, pykF | SNP-4187619-A, DEL-3905639-1, MOB-1907448-IS5-4, SNP-4161966-A, SNP-1352926-T, SNP-3449388-T, MOB-1755687-IS5-4 |
| IBUA2-6 | rpsC, yobF, yijD, bglF, sapD, rpoC, pykF | SNP-4187619-A, DEL-3905639-1, MOB-1907448-IS5-4, INS-3449508-GAACATAACGCGACG, SNP-4161966-A, SNP-1352926-T, MOB-1755687-IS5-4 |
| IBUA2-9 | rpsC, yobF, yijD, bglF, sapD, rpoC, pykF | SNP-4187619-A, DEL-3905639-1, MOB-1907448-IS5-4, SNP-4161966-A, SNP-1352926-T, SNP-3449388-T, MOB-1755687-IS5-4 |
| IBUA3-10 | pykF, sapF, ydbA, rpoB | SNP-4182820-T, SNP-1352163-A, MOB-1472662-IS5-4, MOB-1757082-+G-S5 |
| IBUA4-1 | yjjQ, pykF, sapB, rpoB | SNP-4182820-T, SNP-4603494-A, SNP-1354686-C, INS-1756894-TG |
| IBUA4-8 | rpsD, pykF, rpoB | SNP-3441417-A, SNP-4182820-T, INS-1756894-TG |
| IBUA4-9 | yaiP, speA, infA, pykF, rpoB, bglG | SNP-383289-T, SNP-4182820-T, DEL-3084357-1, SNP-926293-T, SNP-3906597-A, INS-1756894-TG |
| IBUA5-2 | rpoS, pykF, prfA, rpoB | SNP-2866767-T, SNP-4182820-T, SNP-1265009-A, SNP-1756217-T |
| IBUA5-6 | rpoS, pykF, prfA, rpoB | SNP-1265009-A, SNP-1756217-T, SNP-4182820-T, DEL-2867337-96 |
| IBUA6-7 | glyQ, rpoS, pykF, rpoB, infB, pyrE | SNP-3315513-G, DEL-3815808-1, INS-1756495-A, SNP-3725175-G, DEL-2867356-1, SNP-4184792-T |
| IBUA6-9 | glyQ, rne, prfA, rpoC, pykF, ybbW | MOB-538086-IS5-4, INS-1756495-A, SNP-3725175-G, SNP-1143323-T, MOB-4281707-IS5-4, SNP-1265009-A, SNP-4187214-C |
| IBUA7-6 | sapC, pykF, rpoB | SNP-1756434-G, SNP-4182820-T, SNP-1354314-G |
| IBUA7-7 | sapC, pykF, rpsL, rpoB, lysU | SNP-4182820-T, SNP-4354843-T, SNP-1756434-G, SNP-3474485-A, SNP-1354314-G |
| IBUA7-9 | gadE, pykF, sapC, rpoB | SNP-1756434-G, SNP-4182820-T, SNP-1354314-G |
| IBUA8-3 | pykF, glyQ, ilvH | DEL-3815859-82, SNP-87381-T, SNP-3725175-G, INS-1756495-A, DEL-1995819-40006 |
| IBUA8-4 | pykF, glyQ, ilvH | DEL-1995819-40006, SNP-87381-T, SNP-3725175-G, DEL-3815859-82, INS-1756495-A |
| IBUA8-10 | rrsA, glyQ, ilvN, pykF, yffQ | SNP-3725175-G, INS-1756495-A, SNP-2563402-A, SNP-4037067-C, DEL-3815859-82, SNP-3851044-G |
| OCTA1-3 | rpoA, mreB, arpA, sapB, lit | INS-1198505-AATGATGA, MOB-4222091-IS1-9, SNP-3400673-C, SNP-3440923-T, SNP-1354687-A |
| OCTA1-5 | lit, mreB, rpoA, yejO, sapA | MOB-2290201-IS5-4, SNP-1356297-T, SNP-3400673-C, DEL-1198498-8, SNP-3440923-T |
| OCTA1-9 | lit, rpoA, mreB, sapA | SNP-3440923-T, SNP-1356297-T, SNP-3400673-C, DEL-1198498-8 |
| OCTA2-10 | dusB, cydX, rpoC, rlmH, nrdE | SNP-2803042-A, INS-774243-T, SNP-668691-A, MOB-3410273-IS5-4, INS-4186115-TTCCGCTGG |
| OCTA2-14 | dusB, rpoC, rlmH | SNP-668691-A, INS-4186115-TTCCGCTGG, MOB-3410273-IS5-4 |
| OCTA2-16 | dusB, rpoC, rlmH | SNP-668691-A, INS-4186115-TTCCGCTGG, MOB-3410273-IS5-4 |
| OCTA4-9 | rpoA, trkH, mrdB | SNP-665850-T, SNP-4033217-A, SNP-3440923-T |
| OCTA4-10 | rpoA, trkH, mrdB | SNP-665850-T, SNP-4033217-A, SNP-3440923-T |

| | | |
|---|---|---|
| OCTA4-13 | sapD, rpoA, mreB | SNP-3440923-T, SNP-3400300-A, SNP-1353062-A |
| OCTA5-4 | gtrS, rpoC, ydcI, yihQ | MOB-4069461-IS5-4, SNP-1495028-C, SNP-4186605-C, MOB-2469636-IS5-4 |
| OCTA5-8 | recE, ydcI, rpoC, hns, gtrS, yihQ | MOB-4069461-IS5-4, SNP-4186605-C, MOB-1293196-IS5-4, MOB-1416518-IS5-4, SNP-1495028-C, MOB-2469647-IS5-4 |
| OCTA5-9 | rpoC, ydcI, yihQ | MOB-4069461-IS5-4, SNP-4186605-C, SNP-1495028-C |
| OCTA7-2 | dusB, rpoC, mreC, pyrE, ycfQ | DEL-3410240-1, DEL-3815808-1, SNP-1168895-T, SNP-4186605-C, SNP-3399666-A |
| OCTA7-9 | mreC, yfcZ, ycfQ, rpoC, dusB, pyrE | DEL-3815808-1, SNP-4186605-C, SNP-2460805-A, DEL-3410240-1, SNP-1168895-T, SNP-3399666-A |
| OCTA7-10 | dusB, rpoC, mreC, pyrE, ycfQ | DEL-3410240-1, DEL-3815808-1, SNP-1168895-T, SNP-4186605-C, SNP-3399666-A |
| OCTA8-5 | gtrS, hfq | MOB-2469886-IS1-9, SNP-4400417-T |
| OCTA8-7 | gtrS, yciA, hfq | SNP-1311924-C, MOB-2469886-IS1-9, SNP-4400417-T |
| PUTR2-4 | rpoC, cspC, mreB | SNP-4186706-A, INS-1907273-CGTCCTG, SNP-3400986-C |
| PUTR2-6 | rpoC, cspC, mreB | SNP-4186706-A, INS-1907273-CGTCCTG, SNP-3400986-C |
| PUTR3-1 | rph, ygaC, spoT, iscR, lexA, edd, proV, nusG, fliK, icdC | SNP-2661793-A, SNP-2799867-A, INS-2018716-CGGTGGCTG, SNP-3816611-A, SNP-1211308-T, SNP-4257602-T, INS-2804946-T, SNP-3823025-A, SNP-1934806-T, SNP-4178239-T |
| PUTR3-9 | rph, spoT, yphF, yfjW, lexA, pstS, mreB | MOB-2678755-IS5-4, SNP-3816611-A, SNP-3400453-G, SNP-4257602-T, DEL-2774809-1, SNP-3823025-A, INS-3911366-T |
| PUTR3-10 | rph, ygaC, spoT, iscR, lexA, tyrB, proV, nusG, icdC | SNP-2661793-A, SNP-2799867-A, SNP-3816611-A, SNP-4267824-C, SNP-1211308-T, SNP-4257602-T, DEL-2904286-122, INS-2804946-T, SNP-3823025-A, SNP-4178239-T |
| PUTR4-3 | mrdB, cspC, proV, clpX, rpoB | DEL-457406-7, INS-2805131-T, MOB-1907410-IS5-4, SNP-4183154-T, SNP-665554-T |
| PUTR4-7 | mrdB, proV, cspC, rpoB, rpsA | MOB-1907410-IS5-4, INS-2805131-T, SNP-4183154-T, SNP-962473-T, SNP-665554-T |
| PUTR4-8 | glyX, ycgB, proV, rpoB, rpsA, cspC, mrdB | DEL-4392446-1, SNP-4392456-G, SNP-4183154-T, SNP-962473-T, SNP-4392453-T, INS-2805131-T, MOB-1907410-IS5-4, DEL-1236007-50, SNP-665554-T |
| PUTR5-1 | spoT, pykF, fliR, waaS, pstS, mreB, yjhG, ybcK | DEL-3910569-7, SNP-3401016-C, SNP-4522146-A, MOB-2023551-IS5-3, SNP-3823799-T, SNP-568660-T, SNP-1755770-A, DEL-3805056-1 |
| PUTR5-6 | ytfR, rpoC, rpoD | SNP-3214770-C, SNP-4186551-G, SNP-4452005-A |
| PUTR5-8 | rpoC, rpoD | SNP-3214770-C, SNP-4186551-G |
| PUTR6-2 | yieK, stpA, pstA, sspA, murA | DEL-3908805-2, SNP-3899249-G, SNP-3377150-A, SNP-3336073-G, MOB-2798597-IS1-9 |
| PUTR6-7 | yeaR, nmpC, rph, yobF, intE, rpoC, proV, rpsA, cmtB, glnE | MOB-1907448-IS5-4, SNP-4186186-C, DEL-2804864-13, MOB-1879829-Δ1-:, DEL-3815859-82, SNP-3079559-T, SNP-962933-G, SNP-576891-T, DEL-3197154-12, MOB-1199680-IS1-8 |
| PUTR6-10 | yeaR, nmpC, rph, tdcR, yobF, tolA, yjcF, proV, rpsA, cmtB, intE | MOB-1907448-IS5-4, SNP-3267294-T, DEL-777151-48, SNP-4282760-C, DEL-2804864-13, MOB-1879829-Δ1-:, DEL-3815859-82, SNP-3079559-T, SNP-962933-G, SNP-576891-T, MOB-1199680-IS1-8 |
| PUTR7-1 | rpsA, spoT, mreB, nusA | SNP-3823799-A, SNP-3316916-C, SNP-962922-T, SNP-3400811-T |
| PUTR7-7 | rpoD, rpoB, murA | SNP-3214770-C, SNP-3335317-G, SNP-4183154-T |
| PUTR7-9 | rpoD, mdtJ, rpoB, murA | SNP-4183154-T, SNP-3335317-G, DEL-1673532-181, SNP-3214770-C |
| PUTR8-3 | spoT, rpsG, proX, mreB, pyrE, argG | SNP-3400195-A, SNP-3318960-A, DEL-3815808-1, INS-2807248-T, SNP-3473612-C, SNP-3823811-A |
| PUTR8-6 | yedP, spoT, rpsG, nagC, proX, mreB, leuL, pyrE, argG | SNP-2025435-A, SNP-3400195-A, SNP-3318960-A, DEL-3815808-1, INS-2807248-T, DEL-83679-3, DEL-701233-1, SNP-3473612-C, SNP-3823811-A |
| PUTR8-10 | spoT, rpsG, nagC, proX, mreB, sfmH, pyrE, argG | SNP-3400195-A, SNP-3318960-A, DEL-3815808-1, INS-2807248-T, DEL-700785-47, SNP-3473612-C, SNP-562667-C, SNP-3823811-A |

*Supplementary Table 3: Genotypes for the reconstructed strains. Mutation coordinates match the GenBank record for E. coli K-12 MG1655 with accession number NC_000913.*

| STRAIN IDENTIFIER | GENOTYPE |
|---|---|
| **ACRB** | acrB::kan |
| **BIOAB** | bioA/bioB_C809340G |
| **BIOAB_OMPR** | bioA/bioB_C809340G ompR::kan |
| **BIOAB_OMPR_PROQ** | bioA/bioB_C809340G ΔompR proQ::kan |
| **BIOAB_PTRA** | bioA/bioB_C809340G ptrA::kan |
| **BIOAB_PTRA_MDTK** | bioA/bioB_C809340G ΔptrA mdtK::kan |
| **BIOAB_PTRA_OMPR** | bioA/bioB_C809340G ΔptrA ompR::kan |
| **CHER** | cheR::kan |
| **CLSA** | clsA::kan |
| **CSPC** | cspC::kan |
| **CSPC** | cspC::kan |
| **FABR** | fabR::kan |
| **FABR_YFGF** | ΔfabR yfgF::kan |
| **GLYQ** | glyQ_C3725175G |
| **GTRS** | gtrS::kan |
| **ILVH** | ilvH_C87381T |
| **ILVH_GLYQ** | ilvH_C87381T glyQ_C3725175G |
| **ILVN** | ilvN_T3851044G |
| **KGTP** | kgtP::kan |
| **KGTP_PROV** | ΔkgtP proV::kan |
| **KGTP_PROV_YBJL** | ΔkgtP ΔproV ybjL::kan |
| **KGTP_SSPA** | ΔkgtP sspA::kan |
| **KGTP_YBJL** | ΔkgtP ybjL::kan |
| **MARC** | marC::kan |
| **MARC_ROB** | ΔmarC rob::kan |
| **MARC_ROB_MPPA** | ΔmarC Δrob mppA::kan |
| **MARC_ROB_MPPA_YOBF** | ΔmarC Δrob ΔmppA yobF::kan |
| **MARC_ROB_YOBF** | ΔmarC Δrob yobF::kan |
| **MARC_YOBF** | ΔmarC yobF::kan |
| **MDTK** | mdtK::kan |
| **MDTK_OMPR** | ΔmdtK ompR::kan |
| **MDTK_PROQ** | ΔmdtK proQ::kan |
| **MDTK_PTRA** | ΔmdtK ptrA::kan |
| **METJ** | metJ::kan |
| **METJ_ACRB** | ΔmetJ acrB::kan |
| **METJ_PURT** | ΔmetJ purT::kan |
| **METJ_RELA** | ΔmetJ relA::kan |
| **METJ_RELA_ACRB** | ΔmetJ ΔrelA acrB::kan |
| **METJ_RELA_CLSA** | ΔmetJ ΔrelA clsA::kan |
| **METJ_RELA_PURT** | ΔmetJ ΔrelA purT::kan |
| **METJ_RELA_RNB** | ΔmetJ ΔrelA rnb::kan |

| | |
|---|---|
| **METJ_RELA_TREA** | ΔmetJ ΔrelA treA::kan |
| **METJ_RELA_TRER** | ΔmetJ ΔrelA treR::kan |
| **METJ_RELA_YEAR** | ΔmetJ ΔrelA yeaR::kan |
| **MPL** | ΔproV mpl::kan |
| **MPRA** | mprA::kan |
| **MPRA_YHJK** | ΔmprA yhjK::kan |
| **NADR** | nadR::kan |
| **NADR_MPRA** | ΔnadR mprA::kan |
| **NADR_MPRA_YHJK** | ΔnadR ΔmprA yhjK::kan |
| **NADR_RPOA** | ΔnadR rpoA_C3440924G |
| **NADR_YHJK** | ΔnadR yhjK::kan |
| **NUSA_SSPA** | ΔproV nusA_A3317072C sspA_A3377173C |
| **NUSA_SSPA_MPL** | ΔproV sspA_A3377173C nusA_A3317072C mpl::kan |
| **NUSA_SSPA_YBEX** | ΔproV sspA_A3377173C nusA_A3317072C ybeX::kan |
| **NUSA_SSPA_YBEX_MPL** | ΔproV sspA_A3377173C nusA_A3317072C ΔybeX mpl::kan |
| **OMPC** | ompC::kan |
| **OMPR** | ompR::kan |
| **OMPR_OMPC** | ΔompR ompC::kan |
| **OMPR_PROP** | ΔompR proP::kan |
| **OMPR_PROQ** | ΔompR proQ::kan |
| **OMPR_PROQ_MDTK** | ΔompR ΔproQ mdtK::kan |
| **OMPR_PROQ_OMPC** | ΔompR ΔproQ ompC::kan |
| **OMPR_PROQ_PTRA** | ΔompR ΔproQ ptrA::kan |
| **OMPR_PTRA** | ΔompR ptrA::kan |
| **OPGH** | opgH::kan |
| **PHOU** | phoU::kan |
| **PROP** | proP::kan |
| **PROQ** | proQ::kan |
| **PROQ_OMPC** | ΔproQ ompC::kan |
| **PROV** | proV::kan |
| **PROV_ARGG** | ΔproV argG_C3318960A |
| **PROV_CSPC** | ΔproV cspC::kan |
| **PROV_CSPC_MPL** | ΔproV ΔcspC mpl::kan |
| **PROV_CSPC_YBEX** | ΔproV ΔcspC ybeX::kan |
| **PROV_CSPC_YBEX_MPL** | ΔproV ΔcspC ΔybeX mpl::kan |
| **PROV_EDD/ZWF** | ΔproV edd/zwf_C1934806T |
| **PROV_MREB** | ΔproV mreB_G3400195A |
| **PROV_NAGC** | ΔproV nagC::kan |
| **PROV_PTSP** | ΔproV ptsP::kan |
| **PROV_PTSP_MPL** | ΔproV ΔptsP mpl::kan |
| **PROV_PTSP_NAGC** | ΔproV ΔptsP nagC::kan |
| **PROV_PTSP_WBBK** | ΔproV ΔptsP wbbK::kan |
| **PROV_PTSP_YBEX** | ΔproV ΔptsP ybeX::kan |
| **PROV_PTSP_YBEX_MPL** | ΔproV ΔptsP ΔybeX mpl::kan |

| | |
|---|---|
| **PROV_RPSG1** | ΔproV rpsG_A3473612C |
| **PROV_RPSG1_ARGG** | ΔproV rpsG_A3473612C argG_C3318960A |
| **PROV_RPSG1_SPOT** | ΔproV rpsG_A3473612C spoT_G3823811A |
| **PROV_SPOT** | ΔproV spoT_G3823811A |
| **PROV_SSPA** | ΔproV sspA_A3377173C |
| **PROV_WBBK** | ΔproV wbbK::kan |
| **PROV_YGAC_EDD/ZWF** | ΔproV ygaC_C2799867A edd/zwf_C1934806T |
| **PROV_YOBF** | ΔproV yobF::kan |
| **PSTS** | pstS::kan |
| **PTRA** | ptrA::kan |
| **PTSP** | ptsP::kan |
| **PURT** | purT::kan |
| **PYKF** | pykF::kan |
| **PYKF_GLYQ** | ΔpykF glyQ_C3725175G |
| **PYKF_ILVH** | ΔpykF ilvH_C87381T |
| **PYKF_ILVH_GLYQ** | ΔpykF ilvH_C87381T glyQ_C3725175G |
| **PYKF_ILVN** | ΔpykF ilvN_T3851044G |
| **PYKF_RPOS** | ΔpykF rpoS::kan |
| **PYKF_RPOS_YOBF** | ΔpykF ΔrpoS yobF::kan |
| **PYKF_YOBF** | ΔpykF yobF::kan |
| **RELA** | relA::kan |
| **RELA_PURT** | ΔrelA purT::kan |
| **RLMH** | rlmH::kan |
| **ROB** | rob::kan |
| **ROB_YOBF** | Δrob yobF::kan |
| **RPH** | rph::kan |
| **RPOS** | rpoS::kan |
| **RPSG1** | rpsG_A3473612C |
| **RPSG1_MPL** | ΔproV rpsG_A3473612C mpl::kan |
| **RPSG1_MREB** | ΔproV rpsG_A3473612C mreB_G3400195A |
| **RPSG1_MREB_MPL** | ΔproV rpsG_A3473612C mreB_G3400195A mpl::kan |
| **RPSG1_MREB_YBEX** | ΔproV rpsG_A3473612C mreB_G3400195A ybeX::kan |
| **RPSG1_MREB_YBEX_MPL** | ΔproV rpsG_A3473612C mreB_G3400195A ΔybeX mpl::kan |
| **RPSG1_YBEX** | ΔproV rpsG_A3473612C ybeX::kan |
| **RPSG1_YBEX_MPL** | ΔproV rpsG_A3473612C ΔybeX mpl::kan |
| **RZPD** | rzpD::kan |
| **SSPA** | sspA::kan |
| **STFE** | stfE::kan |
| **TREA** | treA::kan |
| **WBBK** | wbbK::kan |
| **YBEX** | ΔproV ybeX::kan |
| **YBJL** | ybjL::kan |
| **YEAR** | yeaR::kan |

| | |
|---|---|
| **YFGF** | yfgF::kan |
| **YGAC** | ΔproV ygaC_C2799867A |
| **YGAC_MPL** | ΔproV ygaC_C2799867A mpl::kan |
| **YGAC_YBEX** | ΔproV ygaC_C2799867A ybeX::kan |
| **YGAC_YBEX_MPL** | ΔproV ygaC_C2799867A ΔybeX mpl::kan |
| **YHJK** | yhjK::kan |
| **YOBF** | yobF::kan |
| **YPJA** | ypjA::kan |
| **YPJA_MPRA** | ΔypjA mprA::kan |
| **YPJA_MPRA_NADR** | ΔypjA ΔmprA nadR::kan |
| **YPJA_MPRA_NADR_YHJK** | ΔypjA ΔmprA ΔnadR yhjK::kan |
| **YPJA_MPRA_YHJK** | ΔypjA ΔmprA yhjK::kan |

*Supplementary Table 4: Mapping of strain identifiers from ALEdb to the strain names used in this study.*

| ALEDB IDENTIFIER | STRAIN NAME |
|---|---|
| **TOL HEXAMETHYLENEDIAMINE A1 F50 I1 R1** | HMDA1-10 |
| **TOL HEXAMETHYLENEDIAMINE A2 F50 I1 R1** | HMDA2-1 |
| **TOL HEXAMETHYLENEDIAMINE A2 F50 I2 R1** | HMDA2-8 |
| **TOL HEXAMETHYLENEDIAMINE A3 F50 I1 R1** | HMDA3-4 |
| **TOL HEXAMETHYLENEDIAMINE A3 F50 I2 R1** | HMDA3-5 |
| **TOL HEXAMETHYLENEDIAMINE A3 F50 I3 R1** | HMDA3-6 |
| **TOL HEXAMETHYLENEDIAMINE A4 F50 I1 R1** | HMDA4-2 |
| **TOL HEXAMETHYLENEDIAMINE A4 F50 I2 R1** | HMDA4-6 |
| **TOL HEXAMETHYLENEDIAMINE A4 F50 I3 R1** | HMDA4-9 |
| **TOL HEXAMETHYLENEDIAMINE A5 F50 I1 R1** | HMDA5-4 |
| **TOL HEXAMETHYLENEDIAMINE A5 F50 I2 R1** | HMDA5-5 |
| **TOL HEXAMETHYLENEDIAMINE A5 F50 I3 R1** | HMDA5-10 |
| **TOL HEXAMETHYLENEDIAMINE A6 F50 I1 R1** | HMDA6-3 |
| **TOL HEXAMETHYLENEDIAMINE A6 F50 I2 R1** | HMDA6-7 |
| **TOL HEXAMETHYLENEDIAMINE A7 F50 I1 R1** | HMDA7-1 |
| **TOL HEXAMETHYLENEDIAMINE A7 F50 I2 R1** | HMDA7-7 |
| **TOL HEXAMETHYLENEDIAMINE A7 F50 I3 R1** | HMDA7-10 |
| **TOL HEXAMETHYLENEDIAMINE A8 F50 I1 R1** | HMDA8-5 |
| **TOL HEXAMETHYLENEDIAMINE A8 F50 I2 R1** | HMDA8-9 |
| **TOL HEXAMETHYLENEDIAMINE A8 F50 I3 R1** | HMDA8-10 |
| **TOL PUTRESCINE A1 F50 I1 R1** | PUTR2-4 |
| **TOL PUTRESCINE A1 F50 I2 R1** | PUTR2-6 |
| **TOL PUTRESCINE A2 F50 I1 R1** | PUTR3-1 |
| **TOL PUTRESCINE A2 F50 I2 R1** | PUTR3-9 |
| **TOL PUTRESCINE A2 F50 I3 R1** | PUTR3-10 |
| **TOL PUTRESCINE A3 F50 I1 R1** | PUTR4-3 |

| | |
|---|---|
| **TOL PUTRESCINE A3 F50 I2 R1** | PUTR4-7 |
| **TOL PUTRESCINE A3 F50 I3 R1** | PUTR4-8 |
| **TOL PUTRESCINE A4 F50 I1 R1** | PUTR5-1 |
| **TOL PUTRESCINE A4 F50 I2 R1** | PUTR5-6 |
| **TOL PUTRESCINE A4 F50 I3 R1** | PUTR5-8 |
| **TOL PUTRESCINE A5 F50 I1 R1** | PUTR6-2 |
| **TOL PUTRESCINE A5 F50 I2 R1** | PUTR6-7 |
| **TOL PUTRESCINE A5 F50 I3 R1** | PUTR6-10 |
| **TOL PUTRESCINE A6 F50 I1 R1** | PUTR7-1 |
| **TOL PUTRESCINE A6 F50 I2 R1** | PUTR7-7 |
| **TOL PUTRESCINE A6 F50 I3 R1** | PUTR7-9 |
| **TOL PUTRESCINE A7 F50 I1 R1** | PUTR8-3 |
| **TOL PUTRESCINE A7 F50 I2 R1** | PUTR8-6 |
| **TOL PUTRESCINE A7 F50 I3 R1** | PUTR8-10 |
| **TOL 1,2-PROPANEDIOL A1 F50 I1 R1** | 12PD1-2 |
| **TOL 1,2-PROPANEDIOL A1 F50 I2 R1** | 12PD1-4 |
| **TOL 1,2-PROPANEDIOL A1 F50 I3 R1** | 12PD1-10 |
| **TOL 1,2-PROPANEDIOL A2 F50 I1 R1** | 12PD2-8 |
| **TOL 1,2-PROPANEDIOL A2 F50 I2 R1** | 12PD2-9 |
| **TOL 1,2-PROPANEDIOL A3 F50 I1 R1** | 12PD3-7 |
| **TOL 1,2-PROPANEDIOL A3 F50 I2 R1** | 12PD3-8 |
| **TOL 1,2-PROPANEDIOL A3 F50 I3 R1** | 12PD3-10 |
| **TOL 1,2-PROPANEDIOL A4 F50 I1 R1** | 12PD4-6 |
| **TOL 1,2-PROPANEDIOL A4 F50 I2 R1** | 12PD4-8 |
| **TOL 1,2-PROPANEDIOL A4 F50 I3 R1** | 12PD4-9 |
| **TOL 1,2-PROPANEDIOL A5 F50 I1 R1** | 12PD5-1 |
| **TOL 1,2-PROPANEDIOL A5 F50 I2 R1** | 12PD5-3 |
| **TOL 1,2-PROPANEDIOL A6 F50 I1 R1** | 12PD6-3 |
| **TOL 1,2-PROPANEDIOL A6 F50 I2 R1** | 12PD6-9 |
| **TOL 1,2-PROPANEDIOL A7 F50 I1 R1** | 12PD7-5 |
| **TOL 1,2-PROPANEDIOL A7 F50 I2 R1** | 12PD7-6 |
| **TOL 1,2-PROPANEDIOL A8 F50 I1 R1** | 12PD8-6 |
| **TOL 1,2-PROPANEDIOL A8 F50 I2 R1** | 12PD8-7 |
| **TOL 1,2-PROPANEDIOL A8 F50 I3 R1** | 12PD8-10 |
| **TOL BUTANEDIOL A1 F50 I1 R1** | 23BD1-6 |
| **TOL BUTANEDIOL A1 F50 I2 R1** | 23BD1-9 |
| **TOL BUTANEDIOL A2 F50 I1 R1** | 23BD2-4 |
| **TOL BUTANEDIOL A2 F50 I2 R1** | 23BD2-7 |
| **TOL BUTANEDIOL A2 F50 I3 R1** | 23BD2-9 |
| **TOL BUTANEDIOL A3 F50 I1 R1** | 23BD3-3 |
| **TOL BUTANEDIOL A3 F50 I2 R1** | 23BD3-4 |

| | |
|---|---|
| **TOL BUTANEDIOL A3 F50 I3 R1** | 23BD3-9 |
| **TOL BUTANEDIOL A4 F50 I1 R1** | 23BD4-3 |
| **TOL BUTANEDIOL A4 F50 I2 R1** | 23BD4-4 |
| **TOL BUTANEDIOL A4 F50 I3 R1** | 23BD4-7 |
| **TOL BUTANEDIOL A5 F50 I1 R1** | 23BD5-1 |
| **TOL BUTANEDIOL A5 F50 I2 R1** | 23BD5-7 |
| **TOL BUTANEDIOL A5 F50 I3 R1** | 23BD5-10 |
| **TOL BUTANEDIOL A6 F50 I1 R1** | 23BD6-1 |
| **TOL BUTANEDIOL A7 F50 I1 R1** | 23BD7-4 |
| **TOL BUTANEDIOL A7 F50 I2 R1** | 23BD7-5 |
| **TOL BUTANEDIOL A7 F50 I3 R1** | 23BD7-7 |
| **TOL BUTANEDIOL A8 F50 I1 R1** | 23BD8-2 |
| **TOL BUTANEDIOL A8 F50 I2 R1** | 23BD8-7 |
| **TOL GLUTARATE A1 F50 I1 R1** | GLUT1-3 |
| **TOL GLUTARATE A1 F50 I2 R1** | GLUT1-9 |
| **TOL GLUTARATE A1 F50 I3 R1** | GLUT1-10 |
| **TOL GLUTARATE A2 F50 I1 R1** | GLUT2-1 |
| **TOL GLUTARATE A2 F50 I2 R1** | GLUT2-9 |
| **TOL GLUTARATE A2 F50 I3 R2** | GLUT2-10-rerun |
| **TOL GLUTARATE A3 F50 I1 R1** | GLUT3-5 |
| **TOL GLUTARATE A3 F50 I2 R2** | GLUT3-7-rerun |
| **TOL GLUTARATE A3 F50 I3 R1** | GLUT3-9 |
| **TOL GLUTARATE A4 F50 I1 R1** | GLUT4-1 |
| **TOL GLUTARATE A4 F50 I2 R2** | GLUT4-4-rerun |
| **TOL GLUTARATE A4 F50 I3 R1** | GLUT4-10 |
| **TOL GLUTARATE A5 F50 I1 R1** | GLUT5-4 |
| **TOL GLUTARATE A5 F50 I2 R1** | GLUT5-5 |
| **TOL GLUTARATE A5 F50 I3 R2** | GLUT5-9-rerun |
| **TOL GLUTARATE A6 F50 I1 R1** | GLUT6-4 |
| **TOL GLUTARATE A6 F50 I2 R2** | GLUT6-5-rerun |
| **TOL GLUTARATE A6 F50 I3 R1** | GLUT6-10 |
| **TOL GLUTARATE A7 F50 I1 R2** | GLUT7-2-rerun |
| **TOL GLUTARATE A7 F50 I2 R1** | GLUT7-6 |
| **TOL GLUTARATE A7 F50 I3 R2** | GLUT7-7-rerun |
| **TOL GLUTARATE A8 F50 I1 R2** | GLUT8-5-rerun |
| **TOL GLUTARATE A8 F50 I2 R1** | GLUT8-6 |
| **TOL GLUTARATE A8 F50 I3 R1** | GLUT8-9 |
| **TOL ADIPIC ACID A1 F50 I1 R1** | ADIP1-1 |
| **TOL ADIPIC ACID A1 F50 I2 R1** | ADIP1-9 |
| **TOL ADIPIC ACID A2 F50 I1 R1** | ADIP2-5 |
| **TOL ADIPIC ACID A2 F50 I2 R1** | ADIP2-6 |

| | |
|---|---|
| **TOL ADIPIC ACID A2 F50 I3 R1** | ADIP2-10 |
| **TOL ADIPIC ACID A3 F50 I1 R1** | ADIP3-2 |
| **TOL ADIPIC ACID A3 F50 I2 R1** | ADIP3-4 |
| **TOL ADIPIC ACID A3 F50 I3 R1** | ADIP3-8 |
| **TOL ADIPIC ACID A4 F50 I1 R1** | ADIP4-1 |
| **TOL ADIPIC ACID A4 F50 I2 R1** | ADIP4-8 |
| **TOL ADIPIC ACID A5 F50 I1 R1** | ADIP5-2 |
| **TOL ADIPIC ACID A5 F50 I2 R1** | ADIP5-6 |
| **TOL ADIPIC ACID A6 F50 I1 R1** | ADIP6-3 |
| **TOL ADIPIC ACID A6 F50 I2 R1** | ADIP6-9 |
| **TOL ADIPIC ACID A6 F50 I3 R1** | ADIP6-10 |
| **TOL ADIPIC ACID A7 F50 I1 R1** | ADIP7-2 |
| **TOL ADIPIC ACID A7 F50 I2 R1** | ADIP7-5 |
| **TOL ADIPIC ACID A8 F50 I1 R1** | ADIP8-3 |
| **TOL ADIPIC ACID A8 F50 I2 R1** | ADIP8-7 |
| **TOL ADIPIC ACID A8 F50 I3 R1** | ADIP8-10 |
| **TOL HEXANOIC ACID A1 F50 I1 R1** | HEXA1-1 |
| **TOL HEXANOIC ACID A1 F50 I2 R1** | HEXA1-4 |
| **TOL HEXANOIC ACID A1 F50 I3 R1** | HEXA1-5 |
| **TOL HEXANOIC ACID A2 F50 I1 R1** | HEXA2-3 |
| **TOL HEXANOIC ACID A2 F50 I2 R1** | HEXA2-9 |
| **TOL HEXANOIC ACID A2 F50 I3 R1** | HEXA2-10 |
| **TOL HEXANOIC ACID A3 F50 I1 R1** | HEXA3-1 |
| **TOL HEXANOIC ACID A3 F50 I2 R1** | HEXA3-7 |
| **TOL HEXANOIC ACID A3 F50 I3 R1** | HEXA3-9 |
| **TOL HEXANOIC ACID A4 F50 I1 R1** | HEXA4-4 |
| **TOL HEXANOIC ACID A4 F50 I2 R1** | HEXA4-7 |
| **TOL HEXANOIC ACID A4 F50 I3 R1** | HEXA4-10 |
| **TOL HEXANOIC ACID A5 F50 I1 R1** | HEXA6-5 |
| **TOL HEXANOIC ACID A5 F50 I2 R1** | HEXA6-6 |
| **TOL HEXANOIC ACID A5 F50 I3 R1** | HEXA6-7 |
| **TOL HEXANOIC ACID A5 F50 I4 R1** | HEXA6-9 |
| **TOL HEXANOIC ACID A6 F50 I1 R1** | HEXA7-2 |
| **TOL HEXANOIC ACID A7 F50 I1 R1** | HEXA8-1 |
| **TOL HEXANOIC ACID A7 F50 I2 R1** | HEXA8-2 |
| **TOL HEXANOIC ACID A7 F50 I3 R1** | HEXA8-5 |
| **TOL OCTANIC ACID A1 F50 I1 R1** | OCTA1-3 |
| **TOL OCTANIC ACID A1 F50 I2 R1** | OCTA1-5 |
| **TOL OCTANIC ACID A1 F50 I3 R1** | OCTA1-9 |
| **TOL OCTANIC ACID A2 F50 I1 R1** | OCTA2-10 |
| **TOL OCTANIC ACID A2 F50 I2 R1** | OCTA2-14 |

| | |
|---|---|
| **TOL OCTANIC ACID A2 F50 I3 R1** | OCTA2-16 |
| **TOL OCTANIC ACID A3 F50 I1 R1** | OCTA4-9 |
| **TOL OCTANIC ACID A3 F50 I2 R1** | OCTA4-10 |
| **TOL OCTANIC ACID A3 F50 I3 R1** | OCTA4-13 |
| **TOL OCTANIC ACID A4 F50 I1 R1** | OCTA5-4 |
| **TOL OCTANIC ACID A4 F50 I2 R1** | OCTA5-8 |
| **TOL OCTANIC ACID A4 F50 I3 R1** | OCTA5-9 |
| **TOL OCTANIC ACID A5 F50 I1 R1** | OCTA6-5 |
| **TOL OCTANIC ACID A5 F50 I2 R1** | OCTA6-6 |
| **TOL OCTANIC ACID A5 F50 I3 R1** | OCTA6-7 |
| **TOL OCTANIC ACID A6 F50 I1 R1** | OCTA7-2 |
| **TOL OCTANIC ACID A6 F50 I2 R1** | OCTA7-9 |
| **TOL OCTANIC ACID A6 F50 I3 R1** | OCTA7-10 |
| **TOL OCTANIC ACID A7 F50 I1 R1** | OCTA8-5 |
| **TOL OCTANIC ACID A7 F50 I2 R1** | OCTA8-7 |
| **TOL COUMARATE A1 F50 I1 R1** | COUM1-2 |
| **TOL COUMARATE A2 F50 I1 R1** | COUM2-3 |
| **TOL COUMARATE A2 F50 I2 R1** | COUM2-4 |
| **TOL COUMARATE A2 F50 I3 R1** | COUM2-7 |
| **TOL COUMARATE A3 F50 I1 R1** | COUM3-1 |
| **TOL COUMARATE A3 F50 I2 R1** | COUM3-9 |
| **TOL COUMARATE A3 F50 I3 R1** | COUM3-10 |
| **TOL COUMARATE A4 F50 I1 R1** | COUM4-2 |
| **TOL COUMARATE A4 F50 I2 R1** | COUM4-5 |
| **TOL COUMARATE A4 F50 I3 R1** | COUM4-10 |
| **TOL COUMARATE A5 F50 I1 R1** | COUM5-3 |
| **TOL COUMARATE A5 F50 I2 R1** | COUM5-5 |
| **TOL COUMARATE A5 F50 I3 R1** | COUM5-8 |
| **TOL COUMARATE A6 F50 I1 R1** | COUM6-2 |
| **TOL COUMARATE A6 F50 I2 R1** | COUM6-5 |
| **TOL COUMARATE A6 F50 I3 R1** | COUM6-9 |
| **TOL COUMARATE A7 F50 I1 R1** | COUM7-5 |
| **TOL COUMARATE A7 F50 I2 R1** | COUM7-6 |
| **TOL COUMARATE A8 F50 I1 R1** | COUM8-1 |
| **TOL COUMARATE A8 F50 I2 R1** | COUM8-6 |
| **TOL ISOBUTYRIC_ACID A1 F50 I1 R1** | IBUA1-7 |
| **TOL ISOBUTYRIC_ACID A1 F50 I2 R1** | IBUA1-9 |
| **TOL ISOBUTYRIC_ACID A2 F50 I1 R1** | IBUA2-1 |
| **TOL ISOBUTYRIC_ACID A2 F50 I2 R1** | IBUA2-6 |
| **TOL ISOBUTYRIC_ACID A2 F50 I3 R1** | IBUA2-9 |
| **TOL ISOBUTYRIC_ACID A3 F50 I1 R1** | IBUA3-2 |

| | |
|---|---|
| **TOL ISOBUTYRIC_ACID A3 F50 I2 R1** | IBUA3-10 |
| **TOL ISOBUTYRIC_ACID A4 F50 I1 R1** | IBUA4-1 |
| **TOL ISOBUTYRIC_ACID A4 F50 I2 R1** | IBUA4-8 |
| **TOL ISOBUTYRIC_ACID A4 F50 I3 R1** | IBUA4-9 |
| **TOL ISOBUTYRIC_ACID A5 F50 I1 R1** | IBUA5-2 |
| **TOL ISOBUTYRIC_ACID A5 F50 I2 R1** | IBUA5-6 |
| **TOL ISOBUTYRIC_ACID A6 F50 I1 R1** | IBUA6-7 |
| **TOL ISOBUTYRIC_ACID A6 F50 I2 R1** | IBUA6-9 |
| **TOL ISOBUTYRIC_ACID A7 F50 I1 R1** | IBUA7-6 |
| **TOL ISOBUTYRIC_ACID A7 F50 I2 R1** | IBUA7-7 |
| **TOL ISOBUTYRIC_ACID A7 F50 I3 R1** | IBUA7-9 |
| **TOL ISOBUTYRIC_ACID A8 F50 I1 R1** | IBUA8-3 |
| **TOL ISOBUTYRIC_ACID A8 F50 I2 R1** | IBUA8-4 |
| **TOL ISOBUTYRIC_ACID A8 F50 I3 R1** | IBUA8-10 |
| **TOL 20C NBUTANOL A1 F50 I1 R1** | BUT1-2 |
| **TOL 20C NBUTANOL A1 F50 I2 R1** | BUT1-3 |
| **TOL 20C NBUTANOL A1 F50 I3 R1** | BUT1-5 |
| **TOL 20C NBUTANOL A2 F50 I1 R1** | BUT2-9 |
| **TOL 20C NBUTANOL A3 F50 I1 R1** | BUT3-3 |
| **TOL 20C NBUTANOL A3 F50 I2 R1** | BUT3-6 |
| **TOL 20C NBUTANOL A3 F50 I3 R1** | BUT3-7 |
| **TOL 20C NBUTANOL A4 F50 I1 R1** | BUT4-4 |
| **TOL 20C NBUTANOL A4 F50 I2 R1** | BUT4-7 |
| **TOL 20C NBUTANOL A4 F50 I3 R1** | BUT4-9 |
| **TOL 20C NBUTANOL A5 F50 I1 R1** | BUT5-2 |
| **TOL 20C NBUTANOL A5 F50 I2 R1** | BUT5-3 |
| **TOL 20C NBUTANOL A6 F50 I1 R1** | BUT6-1 |
| **TOL 20C NBUTANOL A6 F50 I2 R1** | BUT6-3 |
| **TOL 20C NBUTANOL A6 F50 I3 R1** | BUT6-8 |
| **TOL 20C NBUTANOL A7 F50 I1 R1** | BUT7-6 |
| **TOL 20C NBUTANOL A7 F50 I2 R1** | BUT7-7 |
| **TOL 20C NBUTANOL A7 F50 I3 R1** | BUT7-9 |
| **TOL 20C NBUTANOL A8 F50 I1 R1** | BUT9-7 |
| **TOL 20C NBUTANOL A8 F50 I2 R1** | BUT9-10 |

***Supplementary Table 5:*** *Examples of previous efforts to engineer production of the selected chemicals.*

| COMPOUND | AUTHORS | YEAR | TITLE | DOI |
|---|---|---|---|---|
| ISOBUTYRATE | Zhang et al. | 2011 | A Synthetic Metabolic Pathway for Production of the Platform Chemical Isobutyric Acid | 10.1002/cssc.201100045 |
| COUMARATE | Jendresen et al. | 2015 | Highly Active and Specific Tyrosine Ammonia-Lyases from Diverse Origins Enable Enhanced Production of Aromatic | 10.1128/AEM.00405-15 |

| | | | | |
|---|---|---|---|---|
| | | | Compounds in Bacteria and Saccharomyces cerevisiae | |
| ADIPATE | Zhao et al. | 2018 | Metabolic engineering of Escherichia coli for producing adipic acid through the reverse adipate-degradation pathway | 10.1016/j.ymben.2018.04.002 |
| GLUTARATE | Park et al. | 2013 | Metabolic engineering of Escherichia coli for the production of 5-aminovalerate and glutarate as C5 platform chemicals | 10.1016/j.ymben.2012.11.011 |
| 2,3-BUTANEDIOL | Erian et al. | 2018 | Engineered E. coli W enables efficient 2,3-butanediol production from glucose and sugar beet molasses using defined minimal medium as economic basis | 10.1186/s12934-018-1038-0 |
| 1,2-PROPANEDIOL | Nui et al. | 2019 | Metabolic engineering of Escherichia coli for the de novo stereospecific biosynthesis of 1,2-propanediol through lactic acid | 10.1016/j.mec.2018.e00082 |
| PUTRESCINE | Noh et al. | 2017 | Gene Expression Knockdown by Modulating Synthetic Small RNA Expression in Escherichia coli | 10.1016/j.cels.2017.08.016 |
| OCTANOATE | Henritzi et al. | 2018 | An engineered fatty acid synthase combined with a carboxylic acid reductase enables de novo production of 1-octanol in Saccharomyces cerevisiae | 10.1186/s13068-018-1149-1 |
| HEXANOATE | Volker et al. | 2014 | Fermentative production of short-chain fatty acids in Escherichia coli | 10.1099/mic.0.078329-0 |
| BUTANOL | Pontrelli et al. | 2018 | Directed strain evolution restructures metabolism for 1-butanol production in minimal media | 10.1016/j.ymben.2018.08.004 |

***Supplementary Table 6:*** *List of mutated genes for each chemical.*

| COMPOUND | MUTATED GENES |
|---|---|
| **1,2-PROPANEDIOL** | sspA, rpsA, yeaR, rpoA, metJ, relA, ypjA, lrhA, dusA, fabR, yfgF |
| **2,3-BUTANEDIOL** | rpoC, pyrE, rpoB, spoT, rpsA, yeaR, rnb, mprA, iscR, metJ, gtrS, nusG, relA, lon, nanK, zntR, lacZ, elfA, elfD, acrB, essD, fadB, gabP, flu, purT, ydhK, ybhP, ybeT, yhjA, ygaH, uspC, umuD, treR, tolC |
| **HMDA** | rpoC, pyrE, rpoB, sspA, spoT, rpsA, nagC, proV, nagA, rph, rpsG, rnt, nusA, lexA, mdtK, stpA, ptsP, purL, lhr, mpl, cynR, ampC, kup, gatY, ybeX, ygeG, ygbT, xapR, wbbK, pepA, pstB, pnp |
| **PUTRESCINE** | rpoC, pyrE, rpoB, sspA, spoT, rpsA, nagC, proV, yeaR, yobF, rph, pykF, rpsG, nusA, yedP, iscR, lexA, stpA, pstS, cspC, nusG, ytfR, proX, mrdB, yphF, mreB, nmpC, leuL, murA, mdtJ, edd, cmtB, clpX, argG, icdC, intE, glyX, glnE, fliR, fliK, ycgB, ybcK, yjhG, yjcF, yieK, ygaC, yfjW, rpoD, waaS, pstA, tyrB, tolA, tdcR, sfmH |

| | |
|---|---|
| **GLUTARATE** | rpoC, rpoB, sspA, spoT, nagC, proV, yobF, rnb, nagA, mprA, rnt, sapC, kgtP, lit, yfjL, ytfR, polB, proX, yhiL, nohQ, lldR, dkgA, cspE, csiD, hcaD, hofM, insG, greA, fruB, ydfI, wzzE, ybeF, yagU, yfdC, yliE, yiaT, yhfX, yhfA, ygjP, rplM, roxA, rlmI, rclB, ravA, rrlA, rrlC, uvrD, tomB, tdcD, ssuA, sdaC, rspA, rsmC, aceK |
| **ADIPATE** | pyrE, sspA, spoT, hns, nagC, proV, yeaR, nagA, rph, rnt, kgtP, rpoS, pstS, proQ, yhiL, purL, ligA, malQ, mltD, metL, insI1, allD, lacY, icd, idnR, gltP, pdxJ, ycjG, yehD, ydcD, ybjL, yagL, yfgO, yphC, yneO, yicC, uvrB, ubiE, srmB |
| **HEXANOATE** | rpoC, rpoB, hns, rpoA, pykF, sapA, sapB, yedP, mdtK, yfjL, proQ, ompC, ompR, murG, dosP, cydA, emrY, bioB, barA, glxK, opgH, rhmD, ptrA, prpE, rsd, sfmF, rsfS |
| **OCTANOATE** | rpoC, pyrE, hns, rpoA, sapA, sapB, lit, sapD, ycfQ, hfq, gtrS, mrdB, mreB, nrdE, mreC, cydX, dusB, arpA, yfcZ, yejO, ydcI, yciA, yihQ, rlmH, recE, trkH |
| **ISOBUTYRATE** | rpoC, pyrE, rpoB, yobF, pykF, sapB, sapC, sapF, rpoS, sapD, ptsP, lysU, cheR, bglG, bglF, ilvN, ilvH, infA, infB, insA, glyQ, gadE, yedV, ydbA, ybbW, yaiP, yffQ, yjjQ, yijD, rpsL, rpsD, rpsC, rne, rlmE, prfA, rrsA, speA |
| **COUMARATE** | rpoC, pyrE, rpoB, hns, rpoA, rnb, mprA, rpsG, nusA, sapC, sapF, manY, ycfQ, polB, yphF, ypjA, mrdA, ompN, murC, nadR, mgrB, dacA, dcd, epmB, ccmA, atpI, fimC, glnG, focA, fimD, yecT, ydjH, ydiJ, ypjC, yjiP, yhjK, yhgE, yfiN, rho, rhaT, prlF, tufA, thrA |
| **BUTANOL** | pyrE, yobF, sapA, manY, hfq, cspC, leuA, mppA, marC, adeP, otsB, glmU, ycaN, rraA, rob, pheU, tqsA |

### Supplementary Figures

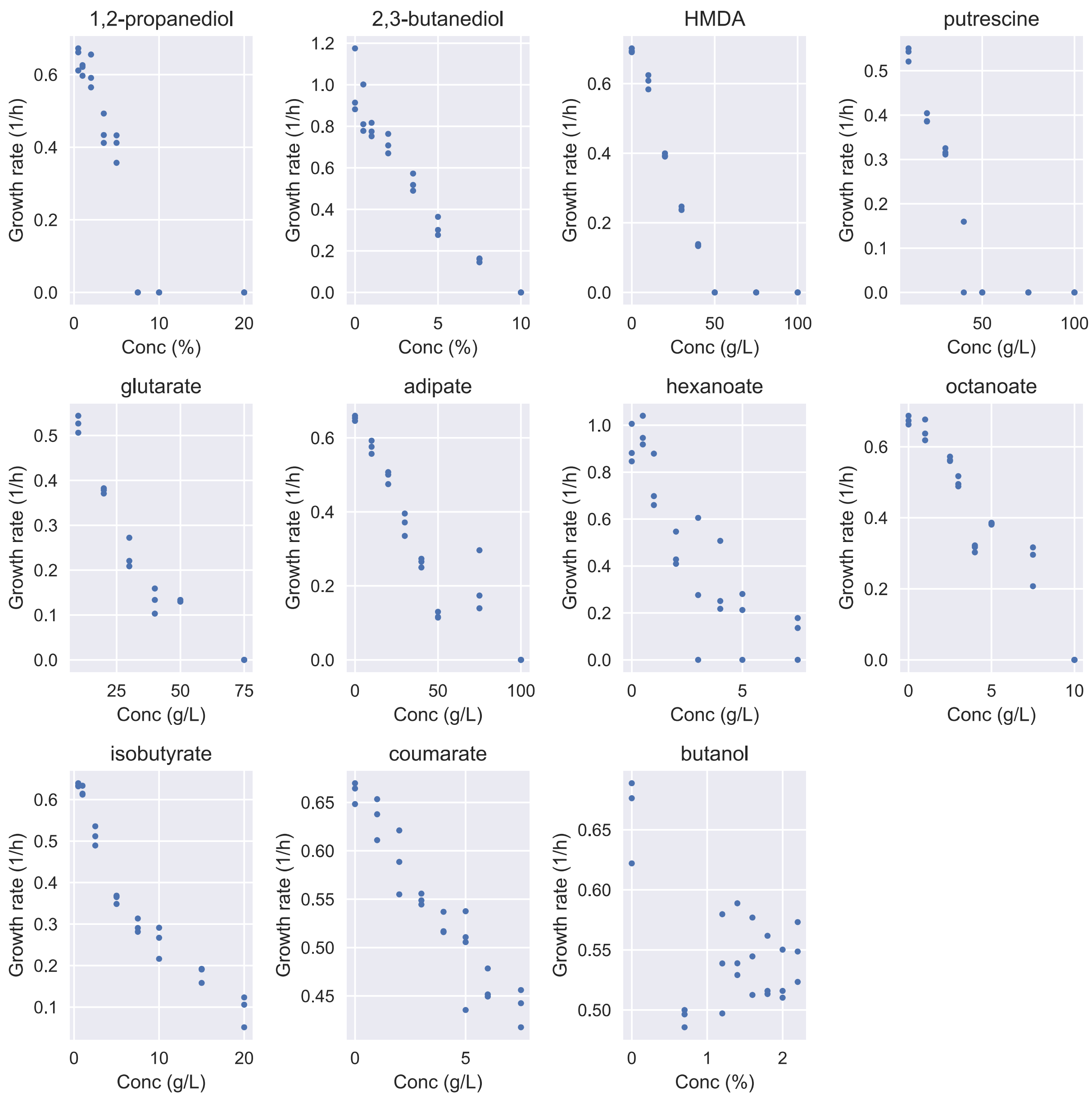


***Supplementary Figure 1:*** *Compound toxicity screening. Growth rates of MG1655 for varying concentrations of the 11 selected compounds. Each individual concentration was tested in biological triplicates.*

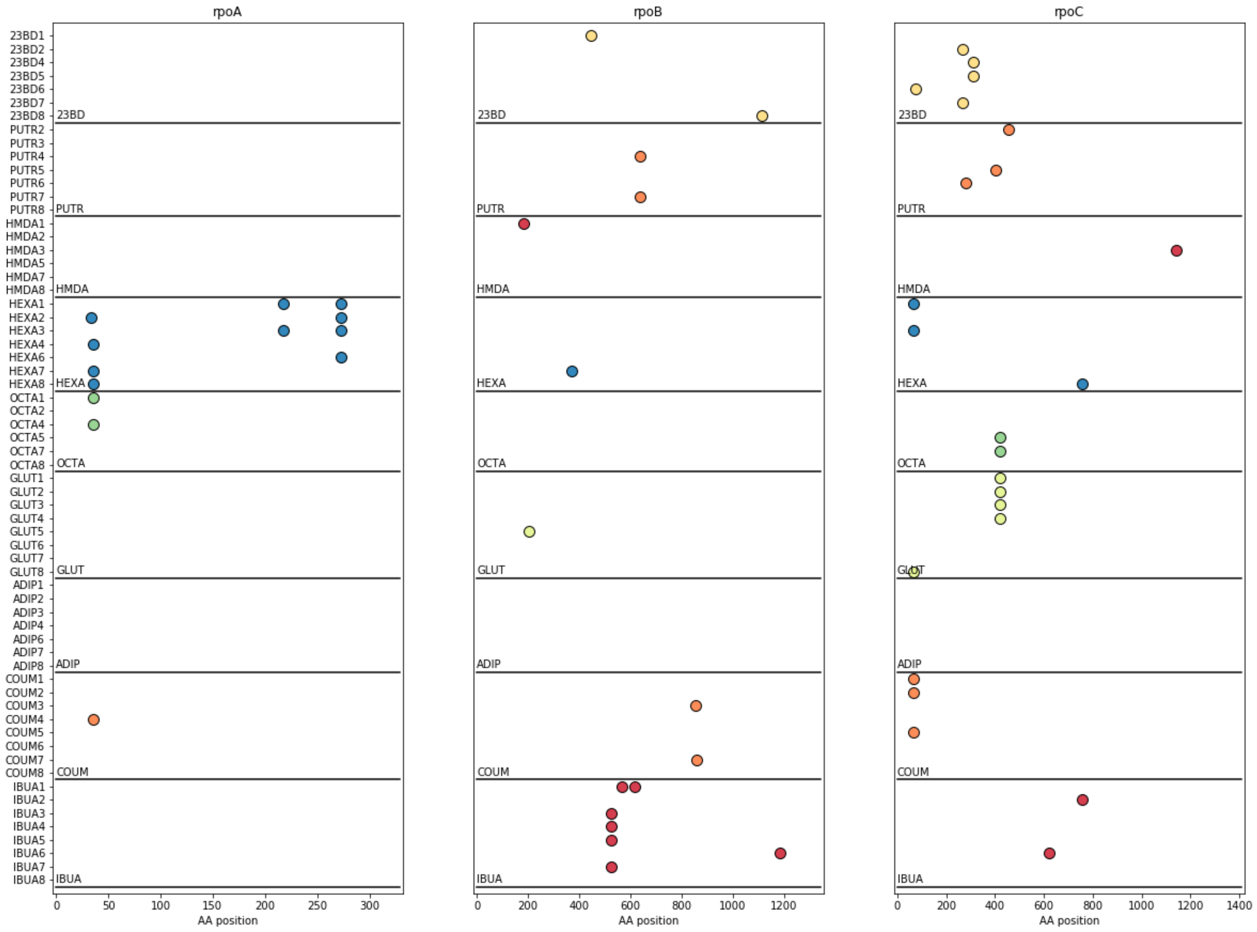


***Supplementary Figure 2:*** *Overview of the locations of observed mutations in RNA polymerase genes. The mutations are shown per population. Mutations found in at least one isolate from a given population are included in the plot.*

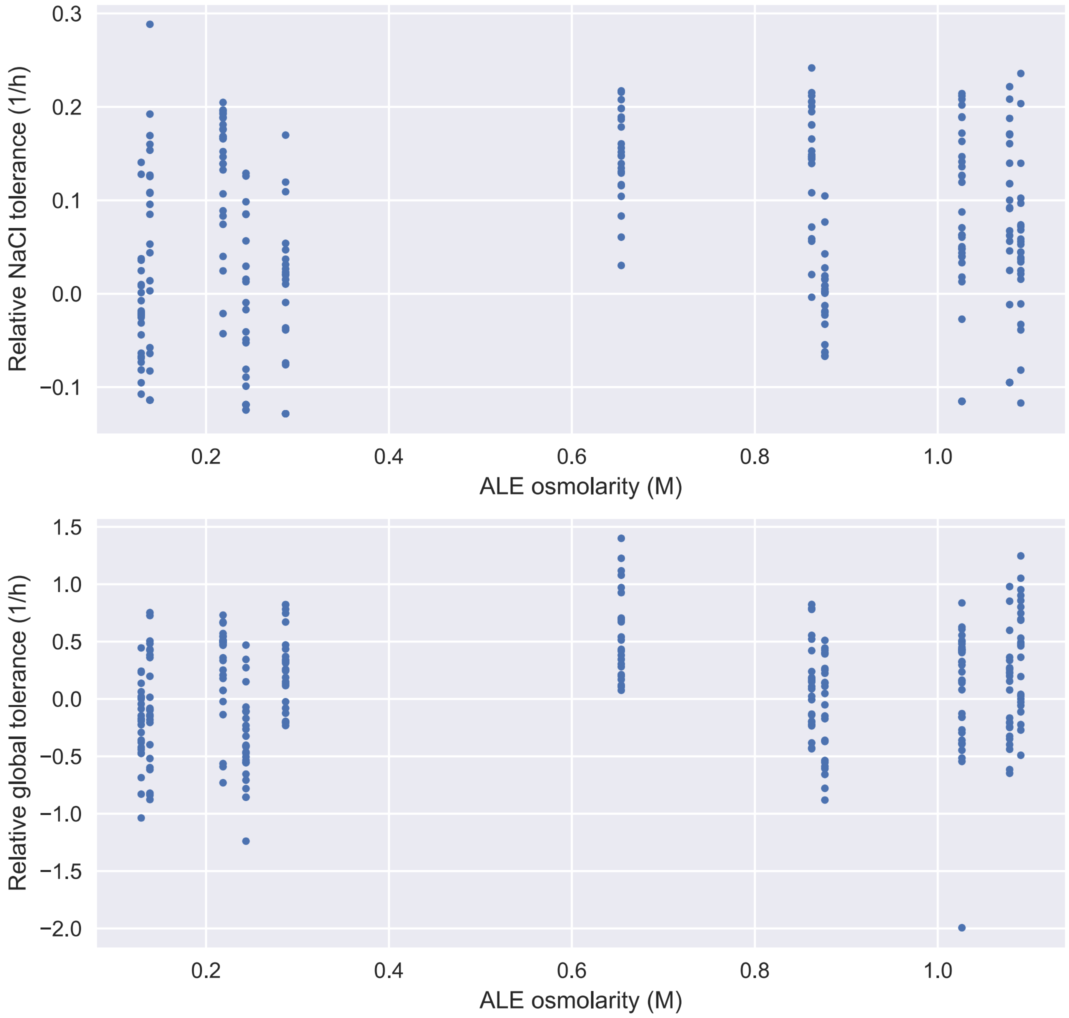


***Supplementary Figure 3:*** *Relationship between final osmolarity of ALE and NaCl tolerance and global tolerance, respectively. Global tolerance is calculated as the mean tolerance to all the 11 chemicals. Each point represents a single evolved strain. The differences in NaCl tolerance and global tolerance between strains from different conditions does not seem to be caused by the differences in osmolarity between the conditions.*

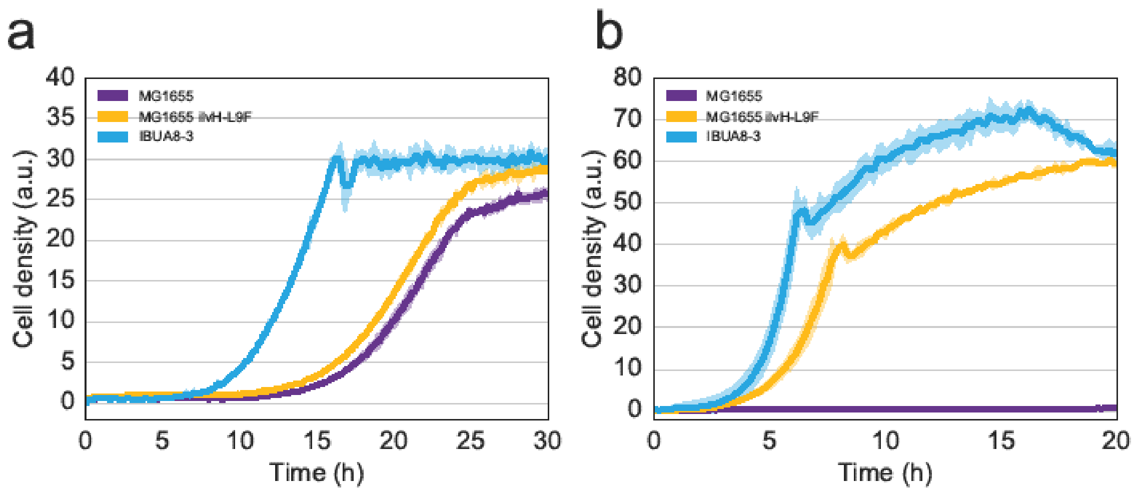


***Supplementary Figure 4****: Isobutyrate tolerance. A) Growth curves in M9 glucose + 10 g/L isobutyrate, for the reference strain, a strain with a L9F mutation in ilvH and the IBUA8-3 evolved strain. b) Growth curves in M9 glucose + 5 g/L valine, for the reference strain, a strain with a L9F mutation in ilvH and the IBUA8-3 evolved strain.*

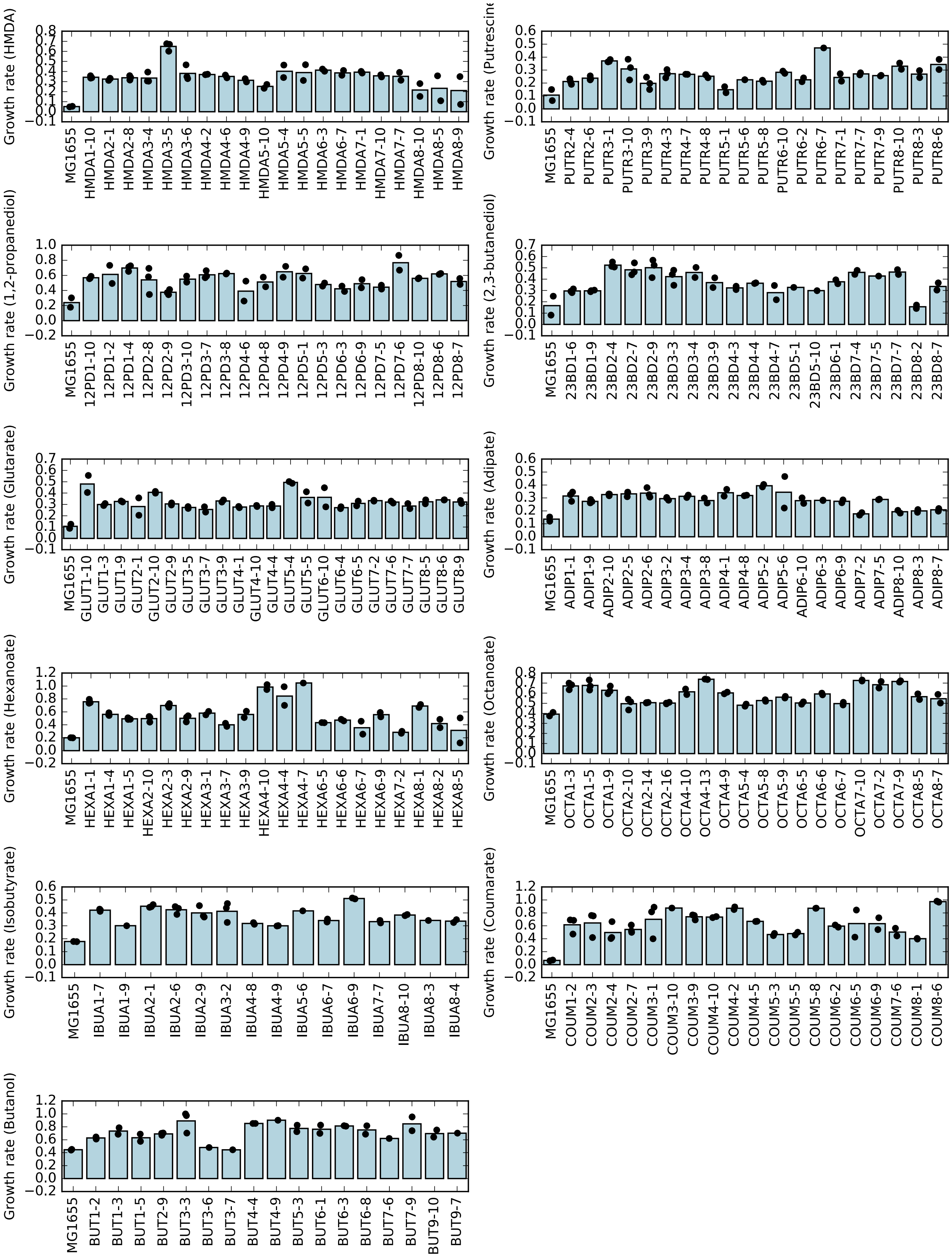


***Supplementary Figure 5:*** *Growth rates of the evolved strains in M9 glucose supplemented with the respective chemical. Chemical concentrations are shown in Supplementary Table 1. Bars represent the mean growth rate, while black dots show the growth rates of each individual biological replicate.*

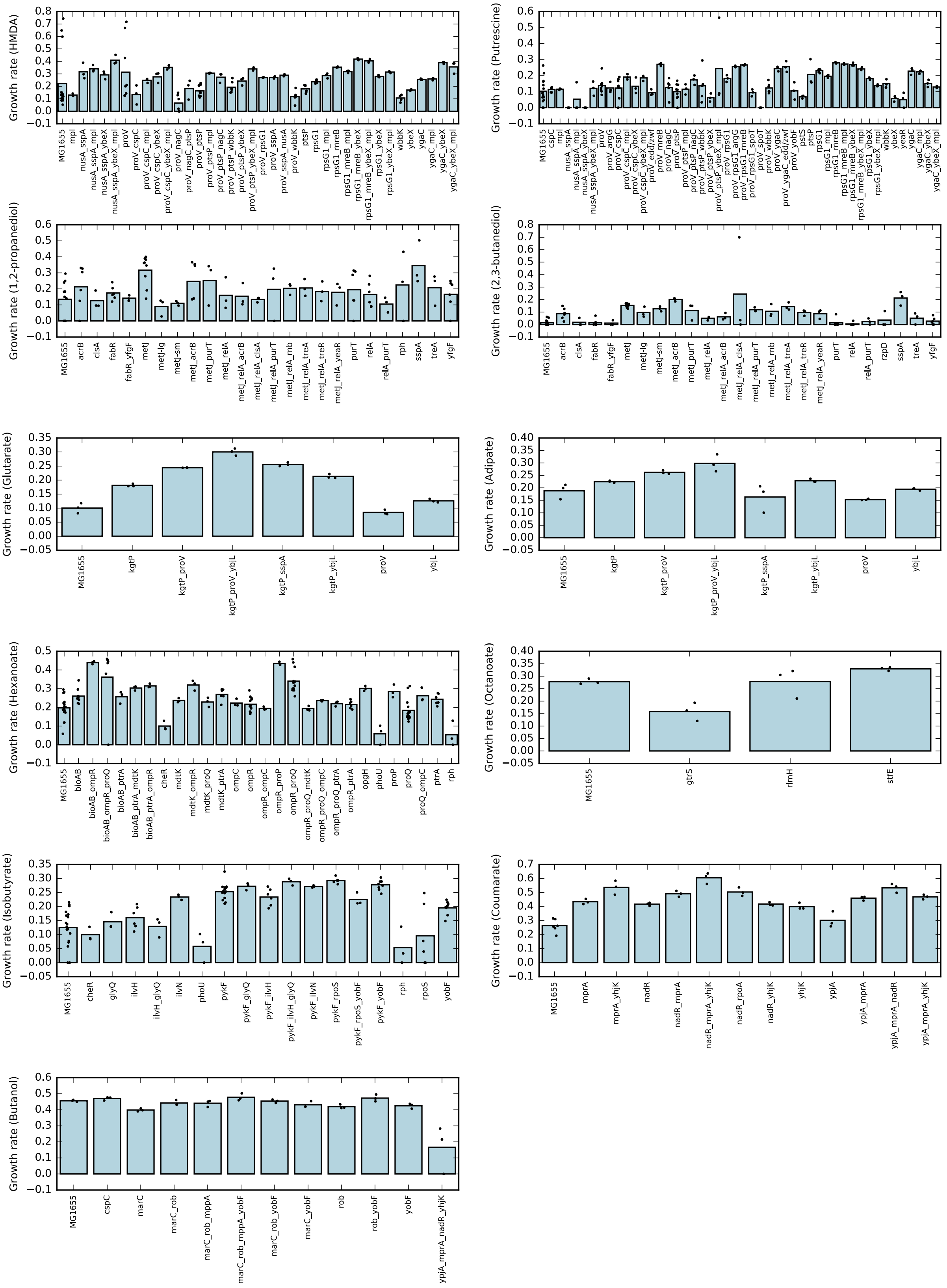
Growth rate (HMDA)
MG1655 mpl nusA_sspA nusA_sspA_mpl nusA_sspA_ybeX nusA_sspA_ybeX_mpl proV proV_cspC proV_cspC_mpl proV_cspC_ybeX proV_cspC_ybeX_mpl proV_nagC proV_nagC_ptsP proV_ptsP proV_ptsP_mpl proV_ptsP_nagC proV_ptsP_wbbK proV_ptsP_ybeX proV_ptsP_ybeX_mpl proV_rpsG1 proV_sspA proV_sspA_nusA proV_wbbK ptsP rpsG1 rpsG1_mpl rpsG1_mreB rpsG1_mreB_mpl rpsG1_mreB_ybeX rpsG1_mreB_ybeX_mpl rpsG1_ybeX rpsG1_ybeX_mpl wbbK ybeX ygaC ygaC_mpl ygaC_ybeX ygaC_ybeX_mpl
Growth rate (Putrescine)
Growth rate (1,2-propanediol)
MG1655 acrB clsA fabR fabR_yfgF metJ metJ-lg metJ-sm metJ_acrB metJ_purT metJ_relA metJ_relA_acrB metJ_relA_clsA metJ_relA_purT metJ_relA_rnb metJ_relA_treA metJ_relA_treR metJ_relA_yeaR purT relA relA_purT rph sspA treA yfgF
Growth rate (2,3-butanediol)
MG1655 acrB clsA fabR fabR_yfgF metJ metJ-lg metJ-sm metJ_acrB metJ_purT metJ_relA metJ_relA_acrB metJ_relA_clsA metJ_relA_purT metJ_relA_rnb metJ_relA_treA metJ_relA_treR metJ_relA_yeaR purT relA relA_purT rzpD sspA treA yfgF
Growth rate (Glutarate)
MG1655 kgtP kgtP_proV kgtP_proV_ybjL kgtP_sspA kgtP_ybjL proV ybjL
Growth rate (Adipate)
MG1655 kgtP kgtP_proV kgtP_proV_ybjL kgtP_sspA kgtP_ybjL proV ybjL
Growth rate (Hexanoate)
MG1655 bioAB bioAB_ompR bioAB_ompR_proQ bioAB_ptrA bioAB_ptrA_mdtK bioAB_ptrA_ompR cheR mdtK mdtK_ompR mdtK_proQ mdtK_ptrA ompC ompR ompR_ompC ompR_proP ompR_proQ ompR_proQ_mdtK ompR_proQ_ompC ompR_proQ_ptrA ompR_ptrA opgH phoU proP proQ proQ_ompC ptrA rph
Growth rate (Octanoate)
MG1655 gtrS rfmH stfE
Growth rate (Isobutyrate)
MG1655 cheR glyQ ilvH ilvH_glyQ ilvN phoU pykF pykF_glyQ pykF_ilvH pykF_ilvH_glyQ pykF_ilvN pykF_rpoS pykF_rpoS_yobF pykF_yobF rph rpoS yobF
Growth rate (Coumarate)
MG1655 mprA mprA_yhjK nadR nadR_mprA nadR_mprA_yhjK nadR_rpoA nadR_yhjK yhjK ypjA ypjA_mprA ypjA_mprA_nadR ypjA_mprA_yhjK
Growth rate (Butanol)
MG1655 cspC marC marC_rob marC_rob_mppA marC_rob_mppA_yobF marC_rob_yobF marC_yobF rob rob_yobF yobF ypjA_mprA_nadR_yhjK

***Supplementary Figure 6:*** *Growth rates of reconstructed strains in M9 glucose supplemented with the respective chemical. Chemical concentrations are shown in Supplementary Table 1. See Supplementary Table 3 for strain genotypes.*
